# Neural and Computational Mechanisms Underlying One-shot Perceptual Learning in Humans

**DOI:** 10.1101/2025.05.09.635727

**Authors:** Ayaka Hachisuka, Jonathan D. Shor, Xujin Chris Liu, Daniel Friedman, Patricia Dugan, Ignacio Saez, Fedor E. Panov, Yao Wang, Werner Doyle, Orrin Devinsky, Eric K. Oermann, Biyu J. He

## Abstract

The ability to quickly learn and generalize is one of the brain’s most impressive feats and recreating it remains a major challenge for modern artificial intelligence research. One of the most mysterious one-shot learning abilities displayed by humans is one-shot perceptual learning, whereby a single viewing experience drastically alters visual perception in a long-lasting manner. Where in the brain one-shot perceptual learning occurs and what mechanisms support it remain enigmatic. Combining psychophysics, 7T fMRI, and intracranial recordings, we identify high-level visual cortex as the most likely neural substrate wherein neural plasticity supports one-shot perceptual learning. We further develop a novel deep neural network model incorporating top-down feedback into a vision transformer, which recapitulates and predicts human behavior. The prior knowledge learnt by this model is highly similar to the neural code in the human high-level visual cortex. These results reveal the neurocomputational mechanisms underlying one-shot perceptual learning in humans.

## INTRODUCTION

The human perceptual system is incredibly malleable even in adulthood. Visual perceptual abilities, from low-level contrast and color sensitivity to high-level expertise of recognizing clinical features in radiological images, can improve dramatically with repeated training^1,2^—“practice makes perfect”. While perceptual learning is often studied in the context of slow, laborious training, it can also occur with a single experience in a drastic, long-lasting manner (an “aha!” moment), a phenomenon termed ‘one-shot perceptual learning’^3–5^. This phenomenon is famously illustrated by the Dalmatian Dog picture^6^ and studied in the laboratory using the “Mooney image” paradigm, wherein degraded images are difficult to recognize initially, but effortlessly recognized once the subject views the corresponding original clear images, and the learning effect lasts many months^3,5,7^. Thus, the visual system possesses very fast learning mechanisms without sacrificing stability or suffering from catastrophic interference. To date, the neural mechanisms underlying this rapid perceptual learning ability remain elusive.

Although artificial intelligence (AI) has shown tremendous progress in basic object recognition over the past decade, one- or few-shot learning remains an unmet need and has emerged as an active area of research in recent years. These research efforts have focused on tasks belonging to concept learning, such as the classification or detection of a novel object based on a few or no training examples^8–12^. Approaches broadly involve learning representations that can be used to distinguish novel cases^12^, learning parameters that can be easily adapted to novel tasks, or learning models of the generating process behind potential novel cases^13^. However, there are several reasons to consider one-shot perceptual learning and one-shot concept learning as fundamentally different phenomena. First, one-shot perceptual learning relies on existing concepts, without the need to form or handle new concepts. Second, existing evidence suggests that they likely rely on different brain structures: the hippocampus and associated medial-temporal lobe structures for concept learning^14,15^, and hippocampus-independent cortical mechanisms for one-shot perceptual learning^4^. Third, very young children can learn and generalize new concepts quickly^16–18^, while one-shot perceptual learning has a protracted developmental time course, reaching adult-level in adolescence^19,20^.

What neural and computational mechanisms support one-shot perceptual learning in humans? Conventional wisdom holds that one-shot, fast learning requires the hippocampus, but a recent study^4^ ruled out this possibility for one-shot *perceptual* learning: memory-impaired patients with bilateral hippocampal lesions were intact at one-shot perceptual learning. This study also demonstrated a clear dissociation between one-shot perceptual learning and episodic memory—both are fast, one-shot learning, but only episodic memory (about whether a picture was previously encountered) is impaired after damage to the hippocampus and associated medial temporal lobe structures.

However, this still leaves a vast hypothesis space for where the learning-related plasticity subserving one-shot perceptual learning might occur in the brain. Previous neuroimaging studies have shown widespread cortical activity changes before vs. after one-shot perceptual learning, from early visual cortex and high-level visual regions to frontoparietal (FPN) and default-mode (DMN) networks^21–25^. In all of these regions, after one-shot learning, neural activity patterns triggered by Mooney images contain more information about the image content and become more similar to the activity patterns triggered by the matching original images (which induced learning). However, not all of these brain regions are necessarily involved in the learning process, and it would be uneconomical for the brain to store multiple copies of prior knowledge (i.e., the knowledge learnt by viewing the corresponding grayscale image). A more efficient solution would be to store the learnt prior knowledge in a particular site or a few interconnected sites, and, once reactivated by a matching visual input (degraded image post learning), it could exert widespread influences on neural processing. In this paper, we aim to investigate where priors are stored, their representational format, and potential computational mechanisms.

Because learning-induced plasticity from synaptic changes is not directly measurable by neuroimaging techniques, the site of prior storage (i.e., where learning/plasticity occurs) has remained unresolved. Previous neuroimaging work hypothesized that either FPN or DMN might encode the prior knowledge learnt in one-shot perceptual learning and send this prior information to visual regions^3,26^. However, this hypothesis was based on observations comparing neural activity driven by the same degraded image input before and after viewing the corresponding original clear image, and the activity differences might reflect a region’s involvement in perceptual processing which can be influenced by priors stored elsewhere.

In other lines of work, previous studies using slow, laborious training paradigms to induce perceptual learning have emphasized plasticity within the visual system^1,2^. And a recent study showed that monkey inferotemporal (IT) cortical neurons are equipped with a multiplexed neural code for object perception and long-term memory, such that familiarity (a form of episodic memory) can be read out from the same neuronal population as perception^27^. However, these studies did not specifically address neural plasticity involved in one-shot perceptual learning, which is distinct from episodic memory^4^ and likely differs from slow, laborious perceptual learning^28^. In sum, the exact brain mechanisms supporting one-shot perceptual learning, including the site of learning-related plasticity, remain unknown.

To pinpoint the site of cortical plasticity and the involved computational mechanisms underlying one-shot perceptual learning in humans, we used several convergent approaches in this study: *First*, using psychophysics, we manipulated the prior-inducing image and assessed its effect on learning. This revealed what kind of information is stored in the prior knowledge encoded in the brain, which was then compared with neural coding properties assayed by 7T fMRI to identify which brain regions have neural coding properties compatible with the information content of prior knowledge. *Second*, using intracranial recordings in neurosurgical patients, we assessed the timing latencies of neural activity changes in different brain regions; brain regions with the earliest prior-driven activity changes are more likely to be the site of prior knowledge storage. *Third*, we built a deep neural network (DNN) capable of one-shot perceptual learning, which both captured the overall magnitude of learning effect and predicted image-specific learning outcomes in humans. We then asked which brain region’s neural code is similar to the prior information learnt by the DNN. The convergent results from these three lines of inquiries point to high-level visual cortex (HLVC) as the site of learning-induced plasticity. Our work further reveals potential computational mechanisms involved in one-shot perceptual learning by developing a DNN model capable of capturing human behavior in this task.

## RESULTS

### Invariance properties of perceptual priors

We first replicated previously observed behavioral effects^22,24,25,29^ using a well-established one-shot perceptual learning paradigm (Fig. 1a). Subjects were instructed to verbally identify the content depicted in the Mooney or grayscale image. In ‘original’ trials, the Mooney images and their matching original grayscale images were presented at the same size, retinal location, and orientation (Fig 1b, top). In ‘catch’ trials, the grayscale image did not match the corresponding Mooney image, which controlled for repetition effects (Fig. 1b, middle). Similar to previous studies, we found robust learning effects in ‘original’ but not ‘catch’ trials. A two-way repeated-measures ANOVA on image recognition rate revealed significant main effects ([Pre vs. Post]: F_1,29_ = 115.5, p < 0.001; [Original vs. Catch]: F_1,29_ = 12.9, p = 0.001) and, critically, a significant interaction effect (F_1,29_ = 39.6, p = 7×10^−7^; for full statistics, see Supplementary Table 1) (Fig. 1d).

**Figure 1:**
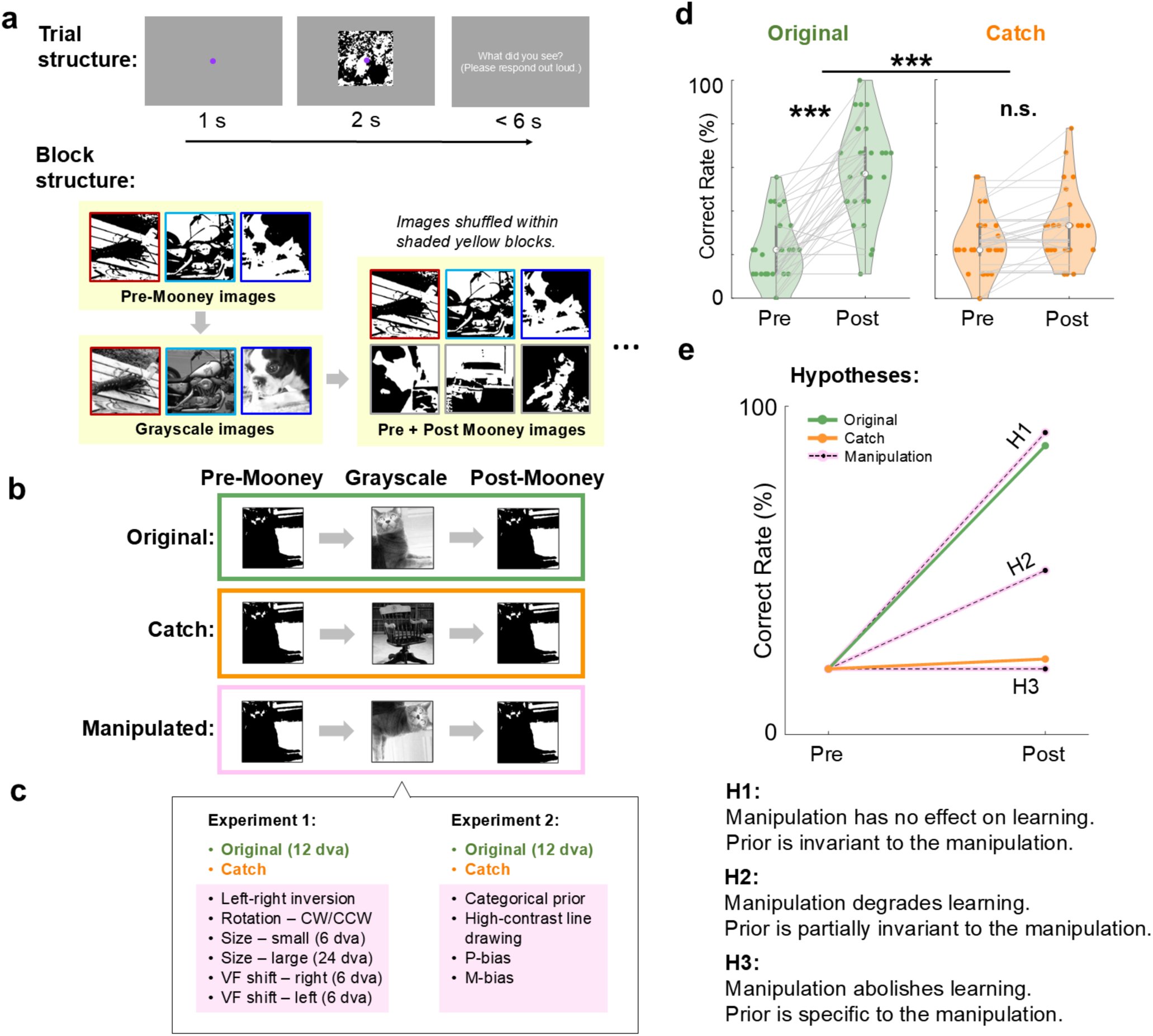
Paradigm and hypotheses for the psychophysics experiment. (**a**) Paradigm. Top: Trial-level timing; images were presented for 2s, followed by a verbal response. Bottom: Block structure; pre and Post Mooney images were shuffled to prevent low-level priming effects. Border colors reflect paired Mooney-grayscale images and were not shown to subjects. (**b**) For each subject, a given Mooney image and its paired grayscale image are presented in one of three conditions (original, catch, manipulated). (**c**) Grayscale image manipulation conditions in each experiment. (**d**) Image identification accuracy for pre- and post-Mooney images in original and catch trials. Data from Experiment 1 (N=30). ***: p < 0.001. (**e**) Hypotheses. Grayscale image manipulation may have no effect on learning (H1), degrade learning without abolishing it (H2), or abolish learning (H3).

To investigate the information content of prior knowledge acquired during one-shot perceptual learning, we manipulated the matching grayscale image (Fig. 1b, bottom) in multiple ways across two experiments (Fig. 1c). We reasoned that if a particular manipulation did not impair learning as compared to the ‘original’ trials (Fig. 1e, H1), it would suggest that the stored perceptual priors are invariant to this manipulation (i.e., did not encode the specific information altered by this manipulation). By contrast, if a particular manipulation abolished the learning effect (Fig. 1e, H3), it would suggest that the perceptual priors are stored in a specific format that the manipulation disrupted. Finally, if a particular manipulation significantly reduced learning but did not abolish it (Fig. 1e, H2), it would suggest that the perceptual priors are partially invariant to that manipulation. Then, the invariance properties of the perceptual priors will indirectly point to where in the brain they are stored given known neural coding properties in different brain regions which we will further validate via an fMRI experiment. We note that our experimental logic is similar to previous psychophysics studies on slow, gradual visual perceptual learning investigating whether the learning effect is specific to the trained condition or transfers to other conditions as a way to shed light on the potential brain loci of learning and plasticity^1,2,30^.

Importantly, to test for one-shot perceptual learning, each Mooney image (presented in both pre and post stages) and its associated grayscale image were presented to a given subject only once under a particular grayscale image condition (original, catch, or a specific manipulation condition; see Fig. 1b). Different images and conditions were presented to different subjects in a counterbalanced design and the results were pooled across unique images and subjects (for details, see SI Methods, Behavioral Experiment 1).

First, to test whether the learnt prior knowledge contains orientation-specific or orientation-invariant information, we left-right inverted the grayscale images or rotated them by 90° (Fig. 2b-c, top). Previous work has shown that orientation-invariant object representations emerge within the primate inferior temporal (IT) cortex^31,32^, where posterior IT is orientation-specific and anterior IT is orientation-invariant^32^, with a similar trend in the human high-level visual cortex (HLVC)^33–35^. We found that both rotation and inversion significantly degraded the learning effect without abolishing it (Fig. 2b-c). A two-way repeated-measures ANOVA comparing each manipulation condition to the ‘original’ trials showed a significant interaction effect ([pre vs. post] × [original vs. manipulated]; inversion: F_1,29_ = 7.4, p = 0.011; rotation: F_1,29_ = 11.2, p = 0.002). Similarly, an ANOVA comparing each condition to the ‘catch’ trials also showed a significant interaction effect (inversion: F_1,29_ = 51.1, p = 7×10^−8^; rotation: F_1,29_ = 20.7, p = 9×10^−5^). Thus, perceptual priors are partially invariant to orientation manipulation.

**Figure 2:**
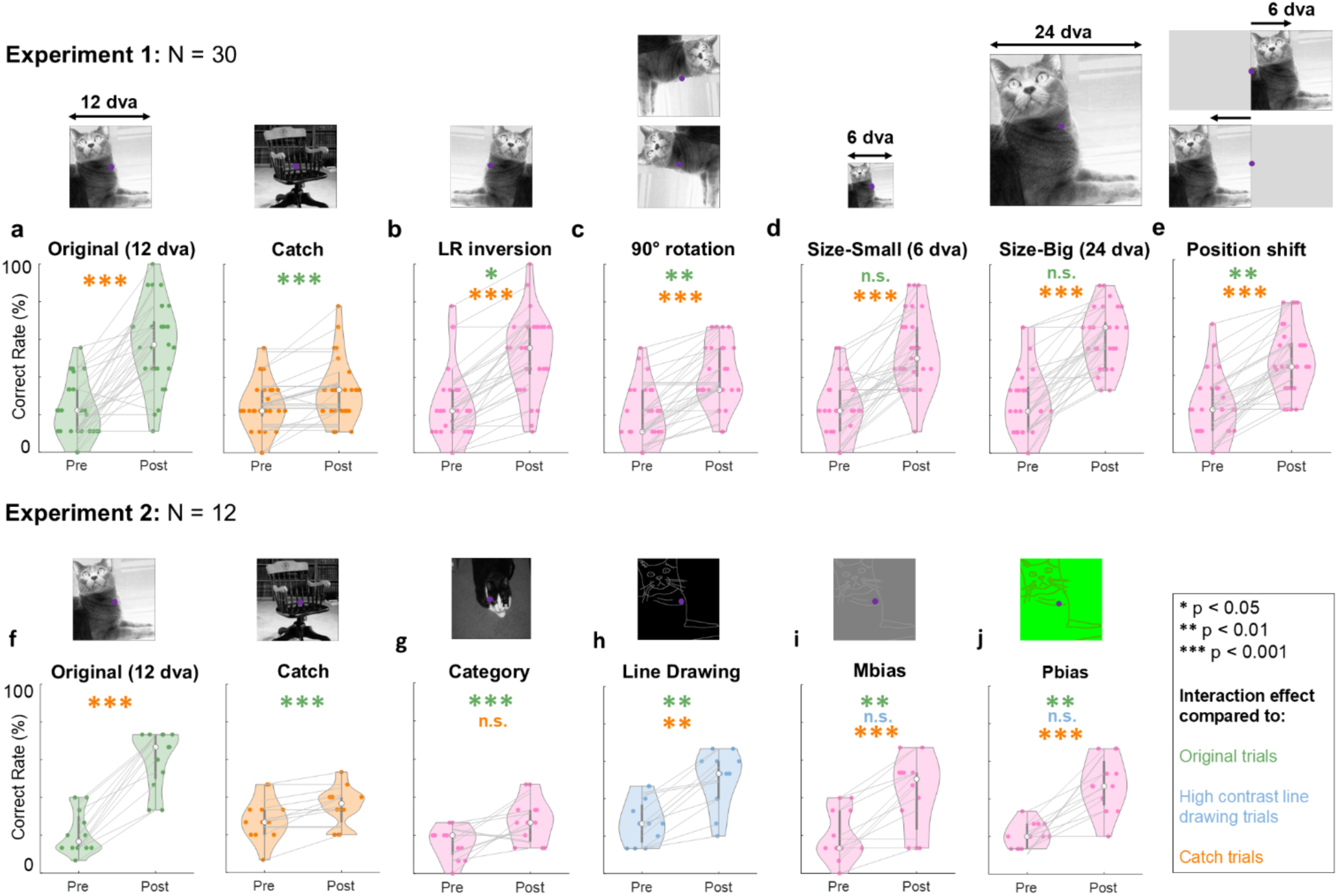
Mapping invariance properties of perceptual priors. Top row: Experiment 1. Bottom row: Experiment 2. (**a** & **f**) Learning effect in response to the original grayscale images and catch images in Experiment 1 and 2, respectively. Data underlying panel **a** are identical to those plotted in Fig. 1d. **(b-e**) Learning effects in response to left-right inverted grayscale images (**b**), 90-degree rotated grayscale images (**c**), size-manipulated grayscale images (**d**), and left/right visual-field shifted grayscale images (**e**). (**g-j**) Learning effects in response to a different grayscale image from the same category (**g**), high-contrast line drawings (**h**), magnocellular pathway-biasing low-contrast images (**i**), and parvocellular pathway-biasing red-green iso-luminant images (**j**). In **i**, image contrast is artificially increased for visualization purpose. Asterisks denote statistically significant interaction effects in a two-way repeated measures ANOVA compared to original (green), high-contrast line drawing (blue), or catch (orange) trials.

Next, we tested for size and positional invariance of the perceptual priors. Given the increasing receptive field size along the visual hierarchy, a given position or size change of the image input may completely alter neuronal encoding in a low-level region while having a modest influence on a higher-level region. Based on previous reports of RF sizes in the ventral visual stream^36,37^, we chose the following size and position manipulations. The original images were presented at central fixation with 12 degrees of visual angle (dva). For size manipulation, we decreased the image size to 6 dva or increased it to 24 dva (Fig. 2d, top). For position manipulation, we shifted the image 6 dva to the left or 6 dva to the right (Fig. 2e, top). A control analysis investigated these manipulations’ impacts on neural coding based on published population receptive field (pRF) data from the human ventral visual stream^37,38^. A voxel’s pRF measures the center and size of its receptive field based on measured fMRI BOLD signal, reflecting an average property across neurons sampled by that voxel. This analysis showed that in anterior HLVC, our chosen size and orientation manipulations have relatively small impacts on neural coding (70–100% pRFs retain diagnostic feature), while position shifts had relatively large impacts (20–40% pRFs) (see Supplementary Fig. 1 and Supplementary Result). In early visual regions, all manipulations have larger impacts on neural encoding (Supplementary Fig. 1).

Strikingly, we found that presenting the grayscale image at double or half the original size had no impact on the learning effect, as shown by non-significant interaction effects compared to the ‘original’ trials (reduced size: F_1,29_ = 1.8, p = 0.189, BF_10_ = 0.7; increased size: F_1,29_ = 0.9; p = 0.354, BF_10_ = 0.4) and significant interaction effects compared to the ‘catch’ trials (reduced size: F_1,29_ = 31.5, p = 5e-6; increased size: F_1,29_ = 45.4; p = 2×10^-7^). Given that size manipulation significantly alters neural coding in early visual cortex (Supplementary Fig. 1c, V1-hV4), these results suggest that the perceptual priors are likely not encoded in the early visual cortex.

We found that position shifts significantly degraded the learning effect yet without completely abolishing it (Fig. 2e), as evidenced by a significant interaction effect as compared to the original trials (F_1,20_ = 8.4, p = 0.007) as well as a significant interaction effect compared to the catch trials (F_1,20_ = 31.2, p = 5×10^-6^). In a control analysis, we excluded trials where subjects shifted their gaze outside a 3-dva radius circle centered on fixation. The results were unchanged with both interaction effects remaining significant (p = 0.031 and p = 0.015; Supplementary Fig. 2).

In sum, orientation manipulations and position shifts significantly impaired learning yet without abolishing it (following H2, Fig. 1e), while size manipulations had no impact on learning (following H1). These results are inconsistent with early visual cortex being a principal site for storing perceptual priors and instead point to HLVC as a likely candidate region. In particular, since orientation invariance emerges within HLVC^32^, if both posterior and anterior HLVC regions are involved in storing the perceptual priors, it would explain the observed pattern of partial invariance to orientation manipulations.

### Perceptual priors are encoded in a perceptual, not conceptual, space

The above experiment used orientation, size and position manipulations to probe invariance properties of the perceptual priors. These can be compared to known neural coding properties along the ventral visual stream where invariance to these manipulations gradually increases across successive stages of neural processing. In a second experiment, we broadened our investigation along two additional lines. First, we probed whether the prior is stored in the perceptual space or at an abstract, conceptual level. To this end, we replaced the grayscale image with another image exemplar from the same object category (Fig. 2g). This manipulation completely abolished learning (following H3, Fig. 1e), as evidenced by a significant interaction effect compared to the original trials (F_1,11_ = 34.4, p = 1×10^-4^, BF_10_ = 3311.7), and a non-significant interaction effect compared to the catch trials (F_1,11_ = 4.29, p = 0.063, BF_10_ = 2.2). This suggests that the perceptual priors are stored in the perceptual space rather than at the conceptual knowledge level, compatible with our hypothesis that it is stored in HLVC, since IT neurons encode category information in a primarily perceptual space with explicit representation of many perceptual features^39,40^.

We further probed whether the magnocellular or parvocellular visual pathway could each support the acquisition of perceptual priors. These two pathways originate from different populations of retinal ganglion cells, and have stronger contributions to the dorsal and ventral visual pathways, respectively, but this separation is not absolute^41,42^. Following previous studies^43–45^, we created line drawings that are either low contrast or red-green iso-luminant, based on the original grayscale images. The low contrast images bias visual processing toward the magnocellular pathway; the red-green iso-luminant image bias visual processing toward the parvocellular pathway. As a control, we created high-contrast line drawing to substitute for the original grayscale images. The high-contrast line drawings induced a significant learning effect that was lower than the original grayscale images (compared to catch: F_1,11_ = 17.0, p = 0.002; compared to original: F_1,11_ = 9.9, p = 0.009), presumably due to the loss of texture and other detailed information. Interestingly, compared to the high-contrast line drawings, neither the M-bias nor the P-bias images caused a significant reduction in the learning effect (M-bias: F_1,11_ = 0.1, p = 0.79, BF_10_ = 0.4; P-bias: F_1,11_ = 0.01, p = 0.91, BF_10_ = 0.4), and both sets of images induced robust learning effects (interaction effect compared to catch, M-bias: F_1,11_ = 24.6, p = 4×10^-4^; P-bias: F_1,11_ = 35.5, p = 1×10^-6^). These results suggest that either the magno- or the parvo-cellular pathway alone can support one-shot perceptual learning. Although the magnocellular pathway has a stronger contribution to the dorsal visual stream, it has collaterals reaching the IT cortex^41^. Therefore, these findings are compatible with our overall hypothesis that the perceptual priors are stored in HLVC.

Finally, a control analysis excluding any trials in which the prior-inducing image was not correctly identified yielded similar results in all conditions of both experiments (Supplementary Fig. 3).

### Neural code in the HLVC matches invariance properties of perceptual priors

To confirm that HLVC indeed has neural coding properties compatible with the information content of the perceptual priors uncovered in our behavioral experiments, we conducted a 7T fMRI experiment using a subset of the grayscale images (Fig. 3a) employed in the behavioral experiments. On each trial, subjects (N=10) viewed a grayscale image presented in the original condition or one of the manipulation conditions employed in Experiment 1 for 500 ms, followed by a 1.5–3 s inter-trial interval.

**Figure 3:**
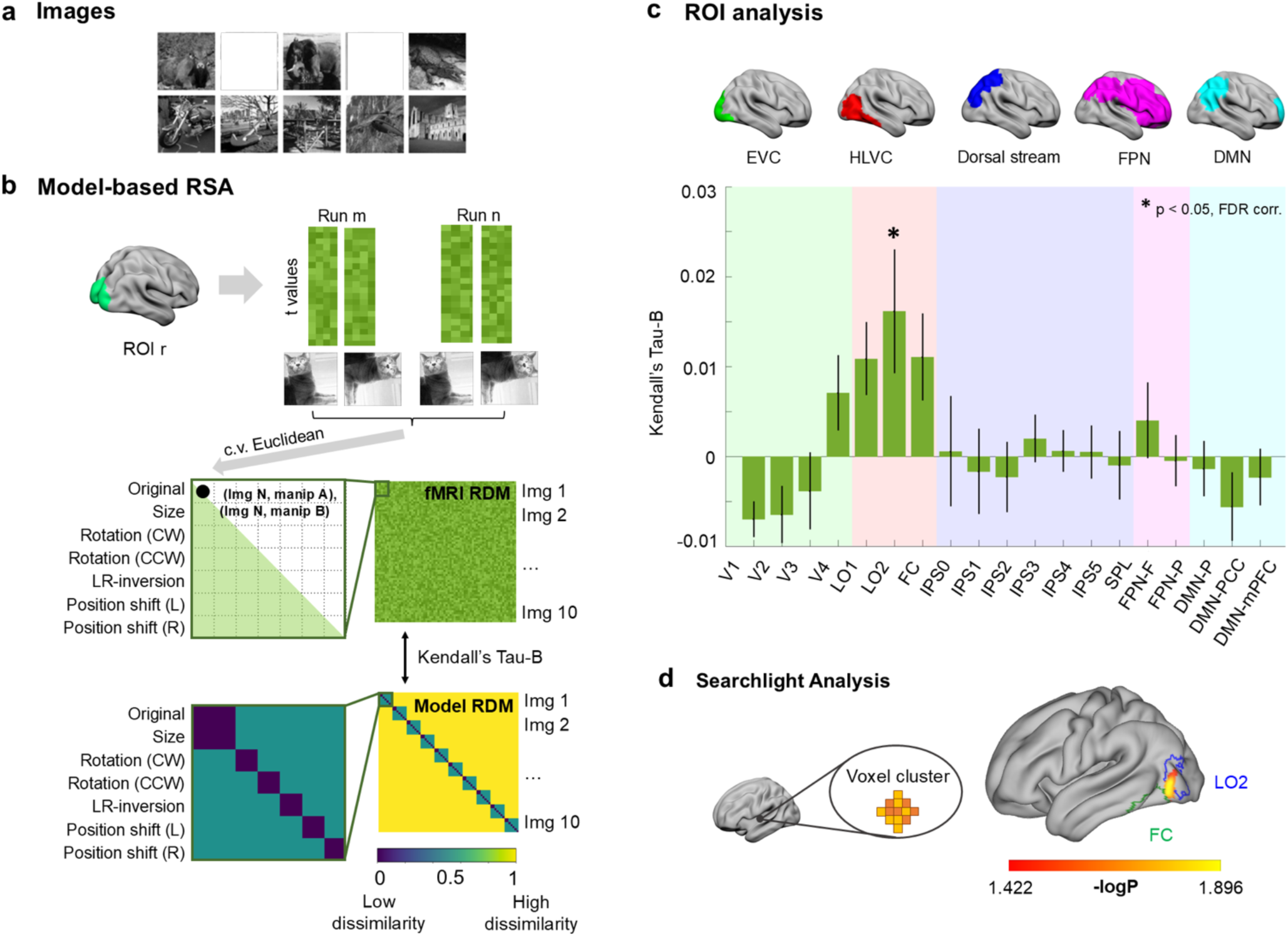
Model-based RSA results (N=10). (**a**) Images used in the fMRI experiment, selected from those used in the psychophysics experiment and evenly distributed between inanimate and animate categories. Images containing people faces have been redacted for privacy concerns. (**b**) Voxel-wise fMRI activation patterns are extracted from each ROI, and cross-validated (c.v.) Euclidean distance was calculated for all pairs of image-condition combination to generate a 70 × 70 RDM. Then, the fMRI RDM is correlated (using Kendall’s Tau-B) with the model RDM that corresponds to behavior results, wherein neural distances are assumed to be high (yellow), intermediate (teal), or low (navy). (**c**) Top: locations of ROIs, grouped by networks; for detailed ROI locations, see Supplementary Fig. 4. Bottom: Kendall’s Tau-B values correlating fMRI and model RDMs, for each ROI in the ventral and dorsal streams, as well as FPN and DMN. Error bars indicate SEM across subjects. For detailed statistics, see Supplementary Table 2. (**d**) A searchlight analysis shows significant correlation between model RDM and fMRI RDM in a voxel cluster within HLVC (*p* < 0.05, cluster-corrected).

For each subject and region of interest (ROI), we computed a neural representational dissimilarity matrix (RDM) comprising of cross-validated (c.v.) Euclidean distances between every pair of image-condition combination computed from voxel-wise fMRI activity patterns. Given 10 unique images and 7 image conditions, this generated a 70 × 70 matrix (Fig. 3b and Supplementary Fig. 5a). ROIs covered early visual cortex (EVC, including V1–V4), HLVC (including LO1, LO2 and FC), as well as FPN and DMN previously shown to be involved in this task^25,26^ (Fig. 3c; for ROI details, see Supplementary Fig. 4 and Methods).

We first tested which ROIs exhibited significant neural invariance to image manipulations. To this end, we averaged within-image, between-condition neural distances (green squares in the RDM shown in Supplementary Fig. 5a; values shown as green bars in Fig. S5b), which were compared against between-image neural distances (sampled from the yellow region of the RDM in Fig. S5a; values shown as yellow ribbon in Fig. S5b; for details, see SI Methods). A significant difference in this test would suggest that the neural representation has significant invariance to image manipulation, since different conditions of the same image are represented more similarly than different images. Significant neural invariance was found in HLVC regions (LO1, LO2, FC) and V4 (all p < 0.01, permutation test, FDR-corrected; black asterisks in Fig. S5b; for full statistics see Supplementary Table 2). A whole-brain searchlight analysis yielded convergent results, with a single significant cluster located within the FC ROI (p < 0.03, cluster-based permutation test; center-of-mass MNI coordinates: [−38, −54, −14], 66 voxels).

We then evaluated how far the neural invariance, when present, is from full invariance in an exploratory analysis. To this end, we compared the within-image between-condition neural distances to zero; a c.v. Euclidean distance of 0 corresponds to what would be expected if there is full invariance (see Methods section “fMRI – Neural Distance Analysis” and Supplementary Methods section “fMRI Analysis—Neural invariance analysis”). Among the four ROIs that contained significant neural invariance to manipulation, only LO2 and FC — the more anterior regions of HLVC — exhibited a non-significant difference from full invariance (Supplementary Fig. 5b, light gray asterisks). This result is consistent with previous work showing that invariant object representation emerges within the IT cortex^32^.

To directly probe neural representations that have similar invariance properties as those identified in our psychophysical experiment for the perceptual priors, we conducted a model-based representational similarity analysis (RSA). We created a model RDM based on the psychophysical results showing that size manipulation had no impact on learning, while orientation and position-shift manipulations significantly degraded the learning effect (Fig. 3b, bottom). Thus, the model RDM contains three levels of neural distance — low (between size manipulation and original), medium (between orientation/position manipulations and original), and high (between different exemplar images). Across all ROIs, model RDM only correlated significantly with neural RDM from HLVC (LO2: p < 0.03, FDR-corrected; Fig 3c). A searchlight analysis across the whole brain also identified a significant cluster within the HLVC (Fig. 3d, p = 0.02, cluster-based permutation test; MNI = [−44, −78, 0], 580 voxels).

Together, these results show that neural representations within the HLVC are uniquely endowed with similar invariance properties as those of the perceptual priors identified by our psychophysical experiment, supporting the notion that HLVC is the prime candidate region for storing the priors in one-shot perceptual learning.

### Learning-induced neural activity changes onset first in the HLVC

The above results show that HLVC is a plausible region for implementing learning-induced plasticity and storing the priors. To further test this hypothesis, we probed the timing properties of neural activity changes induced by one-shot perceptual learning using intracranial EEG (iEEG) recordings in 19 patients undergoing neurosurgical treatment of epilepsy. We reasoned that perceptual priors are stored in latent synaptic connectivity (since one-shot perceptual learning’s effect is long-lasting^4,5^) and, once reactivated by a matching sensory input (e.g., a Mooney image), can trigger widespread changes in neural activity such as those observed in noninvasive neuroimaging^22,24–26^. Therefore, the brain region with the earliest activity changes before vs. after learning is the most likely region for storing the perceptual prior.

In total, 1,886 electrodes were recorded in 19 patients (Fig. 4a; see Supplementary Tables 3 and 4 for demographic, clinical, and electrode information) while patients performed the classic Mooney image task involving ‘original’ and ‘catch’ conditions. Careful screening of patients and collected iEEG data was performed to minimize the potential contribution of pathological activity to the analyzed data (see SI Methods). In all patients, the iEEG electrodes had extensive coverage (Supplementary Fig. 6a) outside the seizure focus (Supplementary Table 4).

**Figure 4.**
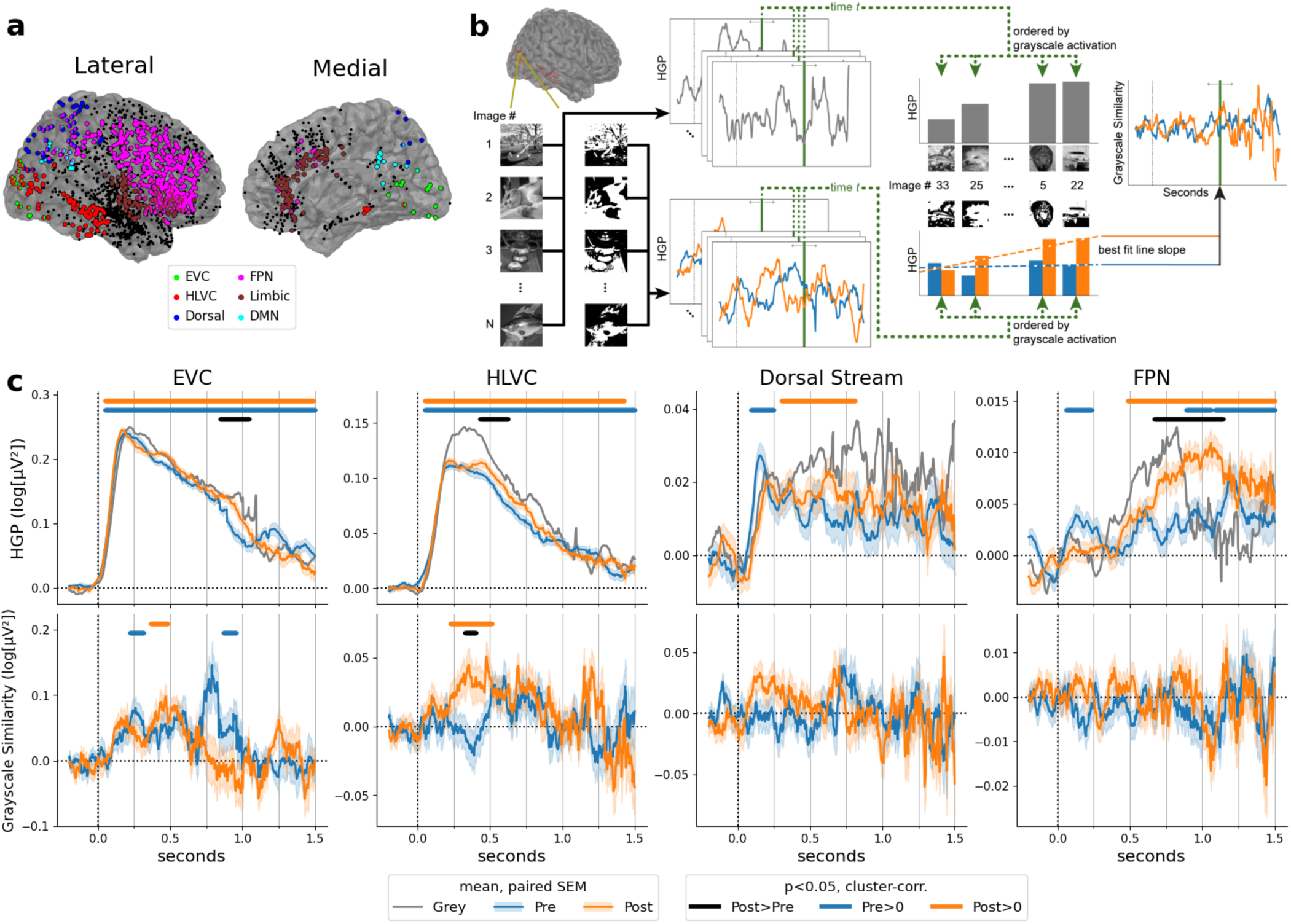
Timing properties of learning-induced activity changes. (**a**) Locations of recorded electrodes from all 19 patients, colored by ROI inclusion. Electrodes in more than one ROIs are correspondingly two-toned. Electrodes not assigned to any ROIs are black. Electrodes in the left hemisphere are shown mirrored across the midline. See Supplementary Table 3 for exact electrode counts by ROI. (**b**) Schematic for Image Preference analysis; for details see Methods. (**c**) Top: results for mean ROI activation analysis, showing the HGP time course in each ROI for each condition. Bottom: results for Image Preference analysis, showing time courses for the neural tuning similarity of pre- and post-images as compared to Grayscale images. Significance bars: p<0.05, cluster-based permutation test (based on Wilcoxon sign-rank test). Shaded areas for the Pre and Post time courses denote SEM corresponding to the paired tests (pre vs. post)^52,53^. See Supplementary Fig. 7 for electrodes included across time course of each ROI.

Patients exhibited similar learning effects as healthy subjects (Supplementary Fig. 6b-c; [pre vs. post] × [original vs. catch]: F_1,18_ = 34.2, p = 1.5 × 10^-5^). In addition to the five networks used in the fMRI analysis (Fig. 3b), we also assessed the limbic network, which included the cingulate, insular and orbitofrontal cortices, given recent results showing the limbic network’s involvement in conscious visual perception^46–48^. Between 32 and 479 electrodes were recorded in each network. In order to maximize the number of trials collected, image presentation ended when a response was given, with a maximal duration of 2 sec (see Methods and Supplementary Fig. 7). For each subject, only images that triggered the classic disambiguation effect—recognized in the post stage and not recognized in the pre stage—entered into the following analysis unless otherwise stated (see SI Methods for details).

We first assessed neural activation time courses, as indexed by high gamma (50–120 Hz) power (HGP)^49,50^, for each perceptual stage (Pre, Grayscale, Post). Both early visual cortex (EVC) and HLVC activated early (*p* < 0.05, cluster-based permutation test), at approximately 50 ms after image onset for both pre- and post-Mooney images (Fig. 4c, top, blue and orange bars). Importantly, post-Mooney images elicited significantly higher neural activity than pre-Mooney images (Fig. 4c, top, black bars) in the HLVC at approximately 430–623 ms (onset time 95% confidence interval [CI]: 191–551 ms), followed by FPN at approximately 668–1,143 ms (onset time 95% CI: 609–943 ms), and later in EVC at approximately 844–1045 ms (onset time 95% CI: 744–893 ms).

Dorsal stream, FPN and DMN all had a relatively early and transient neural activation for pre-Mooney images (significant clusters found at 92–250 ms, 57–234 ms, and 49–186 ms respectively; Fig. 4c and Supplementary Fig. 8, top, blue bars). This early and transient neural activation to pre-Mooney images (<250 ms) was likely trigged by bottom-up visual activation that subsided quickly when recognition was unsuccessful. In addition, FPN had higher neural activity to post-than pre-Mooney images at approximately 668–1,143 ms, which may be related to recognition-triggered decision-related activity. We did not observe significant neural activation to pre- or post-Mooney images in limbic regions.

To pinpoint neural activity specifically related to prior-guided perceptual processing, we followed an earlier approach^26^ to identify time points at which the pre- or post-Mooney image-elicited activity has a similar neural tuning profile as neural activity triggered by the grayscale image (i.e., if an electrode is tuned towards a certain grayscale image, it also exhibits high activity to the matching Mooney image; for analysis schematic, see Fig. 4b). Prior-induced neural activity would manifest as higher neural similarity between post and grayscale images than between pre and grayscale images.

In HLVC, we found that post-Mooney images elicited similar neural tuning profiles as grayscale images at approximately 225–516 ms (Fig. 4c, bottom, orange bar; onset time 95% CI: 152–420 ms), and this similarity is significantly higher than the pre-grayscale similarity (black bar). Importantly, this effect in HLVC preceded that in EVC (at 365–483 ms; Fig. 4c, bottom, orange bar; onset time 95% CI: 242–455 ms), and EVC did not exhibit a significant post vs. pre difference. This result, showing earlier and stronger prior-guided neural activity in HLVC, suggests that feedback from HLVC to EVC could have carried prior-related information. EVC also had two time clusters in which pre-Mooney images had similar neural tuning as grayscale images (Fig. 4c, bottom, blue bars), which can be explained by similar visual features between Mooney images and their matching grayscale images, such as co-localized contours. We did not observe similar neural tuning between pre/post-Mooney images and grayscale images in any other networks (Fig. 4c and Supplementary Fig. 8, bottom).

To further test the idea that the similar neural tuning in HLVC between disambiguated post-Mooney images and their matching grayscale images reflect the influence of one-shot perceptual learning, we performed a control analysis using pre-Mooney images that were spontaneously and correctly recognized before seeing the matching grayscale images. Recognizing the Mooney image prior to viewing the greyscale original version occurs in a minority of trials (Supplementary Fig. 6c) and indicates an alternative source of prior knowledge derived from lifelong experiences, distinct from the one-shot priors acquired by viewing the original greyscale images. We found that spontaneously recognized pre-Mooney images did elicit similar neural tuning profiles as greyscale images, but with a distinct temporal profile to that of the disambiguated post-Mooney images described above (Supplementary Fig. 9). Recognized pre-Mooney images exhibited similarity for two short-lived periods at approximately 262-345 ms and 641-740 ms — possibly related to a feedforward and a feedback wave^51^; by contrast, disambiguated post-Mooney images exhibited similarity in one cluster at approximately 225–516 ms. The broader temporal cluster with a later peak for disambiguated post-Mooney images (at 355 ms as compared to 301 ms) suggests that additional processing is required in HLVC to bring the recently learned prior knowledge to bear as compared to basic object recognition guided by lifelong knowledge.

Together, these results show that prior-induced neural activity onsets first in HLVC (at approximately 225 ms), preceding that in EVC. Strikingly, we did not find prior-induced neural activity in the dorsal visual stream, FPN, DMN or limbic network. This result, obtained from extensive iEEG sampling across cortical networks, provides strong evidence that the perceptual priors are stored and reactivated in HLVC.

### A top-down transformer captures human behavior during the one-shot perceptual learning task

To shed light on potential computational mechanisms underlying one-shot perceptual learning in humans, we sought to develop an image-computable DNN model that can recapitulate human behavior on this task. Instead of modeling specific brain regions, we optimized the model to match human performance, thus avoiding circularity when using the model to localize prior representations in the brain. To preview, we constructed a DNN model which, given a sequence of images, stores accumulated information in a prior module and uses it to modulate visual information processing. We show that our DNN model achieves one-shot perceptual learning capability similar to that of humans, has similar error pattern as human subjects and can be used to predict human learning outcome for a specific image, thereby proving its efficacy to approximate perceptual priors learnt by human subjects. We further show that the prior information learnt by the model has the highest correspondence to neural representation in the human HLVC.

We converted the Mooney image learning task to a computational benchmark to recreate our experimental setup *in silico.* Using this benchmark, we developed a top-down transformer architecture engineered to solely rely on top-down signaling for one-shot learning^54^ (Fig. 5a, see Methods and Supplementary Fig. 10 for details). There are two main components in our model. The first component is a vision backbone (using the transformer architecture), which is pre-trained using self-supervised learning. The second component, key to recapitulating the one-shot learning behavior, is a prior storage module that is responsible for storing prior knowledge about the images seen. Using these two components, we design two pathways for computing the visual representations suitable for the one-shot perceptual learning task: the bottom-up pathway, and the top-down pathway. When an image is first presented to the bottom-up pathway, the vision backbone produces visual representations that are unmodified by previous experiences. The output of the bottom-up pathway is not directly involved in the decision-making process but used as a query to **retrieve** relevant representations from the prior storage module. The relevant context from the prior storage module is then used as **top-down conditioning** to modulate the model in the top-down pathway. Here, the same vision backbone computes image features of the currently shown image again, but this time with the conditioning provided by the prior storage module. Finally, the output of the modulated computation is used to obtain a classification label and **update** the prior module to incorporate the current information.

**Figure 5:**
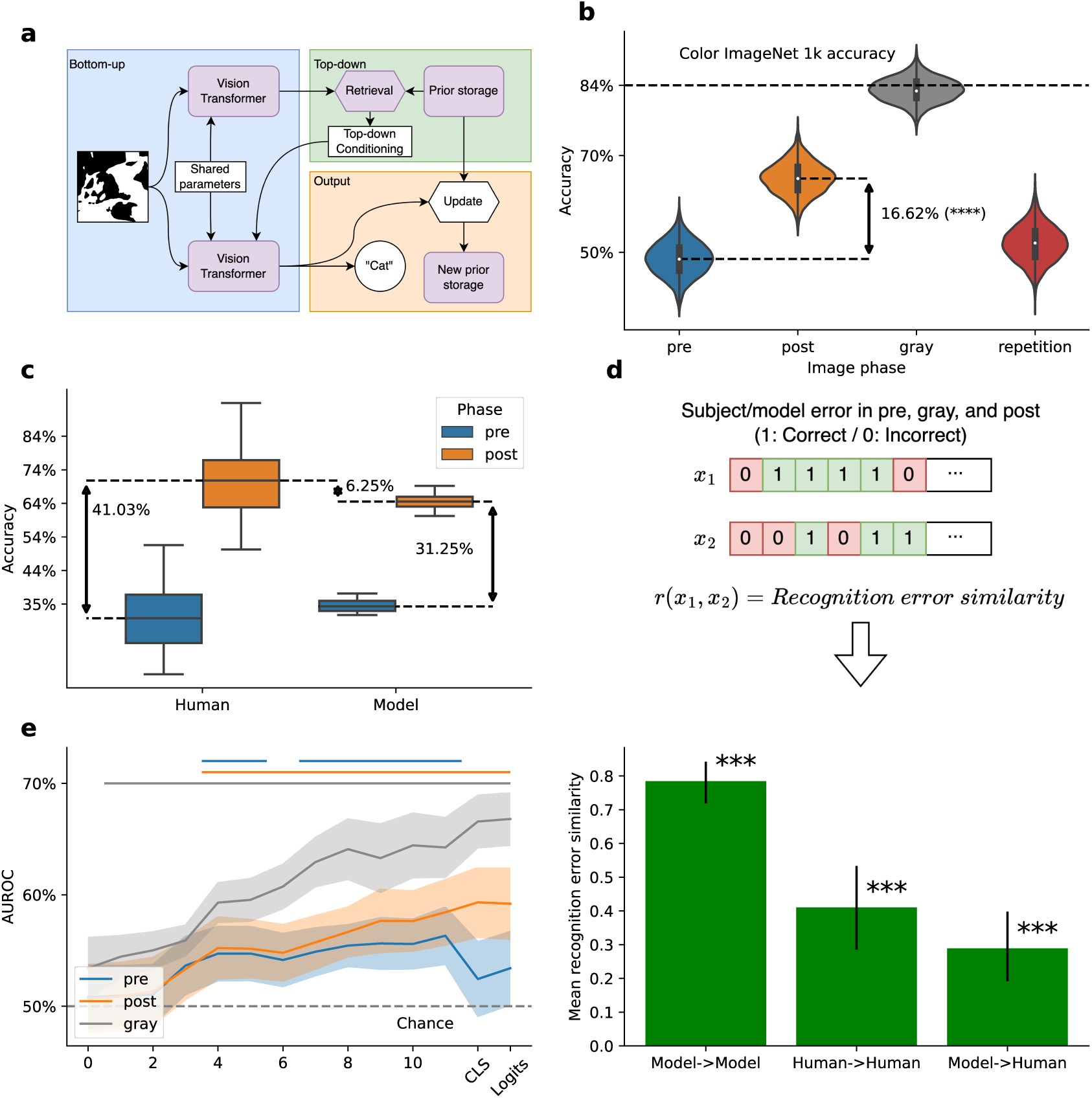
Model displays perceptual learning effect and predicts human learning outcomes. (**a**) DNN model schematic. Model compares bottom-up features with ‘state’ representing prior knowledge, produces top-down conditioning feature, and then produces a final output which is then used to update the model state. For details, see Methods and Supplementary Fig. 10. (**b**) Model accuracy on 1,000 test sequences (sequence length: 630, including 210 unique Mooney images) constructed from grayscale ImageNet 1k images and their Mooney image counterparts. Repetition effect was evaluated on the same Mooney images presented twice in a sequence without matching grayscale images (sequence length: 420). (**c**) Model learning performance as compared to human subjects, where the model was presented with identical image sequences as human subjects. Whiskers show min and max of accuracy. (**d**) Image recognition error pattern similarity between human subjects, between model fed with difference sequences of the same images, and between model and human subjects. Error bars indicate 95% CI of Pearson’s r. (**e**) Human learning outcome prediction. Horizontal bars above show significant above-chance prediction. On x axis, numbers refer to model layer and CLS, Logits refer to model’s representation following its last layer. Individual image learning outcomes can be significantly predicted using the model’s grayscale image representation starting from the second layer onward (p<0.01, max-t permutation test). Shaded areas show 95% CI across subjects.

We first evaluated the model’s performance and perceptual learning effect on 1,000 image sequences generated from randomly chosen grayscale images from ImageNet 1k dataset and their Mooney image counterparts (automatically generated, see Methods), following the same task structure as the human psychophysics study (see block structure in Fig. 1a, without any manipulated grayscale images). We define one-shot perceptual learning effect here as the increase in accuracy in the post-Mooney phase compared to the pre-Mooney phase. Our top-down transformer model displayed an average perceptual learning effect of 16.62% (post – pre; Fig. 5b; pre vs. post, Mann-Whitney U test: p < 0.005, N=1000). This increase is much higher than mere repetition-induced learning effect of 3.11% (Fig. 5b; post vs. repetition, Mann-Whitney U test: p < 0.005, N=1000), indicating that the model exhibits genuine one-shot perceptual learning.

To further evaluate the model’s ability for one-shot perceptual learning against humans, we conducted an online behavioral study (N=12) using a larger set of Mooney images (n=219) (see SI Methods). The 90 images on which human subjects showed the greatest degree of perceptual learning were chosen for the in-person human psychophysics experiments described earlier. We exposed the model to the identical task and image sequences as the human subjects, and plotted model performance against human performance for the top 90 images. Overall, evaluated on the same identical task, the model exhibited a similar perceptual learning effect as human subjects (Fig. 5c), with the absolute post-phase human accuracy at 72% compared to the model’s at 66%.

We compared our novel top-down transformer model with existing well-known neurobiologically motivated DNNs (henceforth ‘baseline models’), including BLT^55^ and CORnet^56^. Our model significantly outperformed these baseline models on the one-shot perceptual learning task, as shown by model performance in the evaluation phase (Supplementary Fig. 11a). In addition, when exposed to the image sequences used in the human online psychophysics experiment, CORnet and BLT had sharply degraded performance and failed to maintain the learning effect (Supplementary Fig. 11b). This drop in performance for baseline models was likely due to their inability to maintain the long-term storage of visual priors (the psychophysics task had much longer image sequences than those used in the model training/evaluation phase).

To confirm that the top-down conditioning from the prior storage module is key to our model’s success, we corrupted this conditioning signal by using weighted average of the conditioning tokens and norm-matched Gaussian noise. As the weight of the noise increases from 0.05 to 0.8, the model’s performance improvement from the pre to post stage drops sharply (Supplementary Fig. 13), confirming that the conditioning by the prior storage module is key to the model’s success at one-shot perceptual learning.

To examine whether our novel DNN model exhibits any behavioral alignment to humans beyond mimicking the overall accuracy, we analyzed the error patterns of human subjects and our model when presented with the same image sequences. The twelve human subjects were each presented with a unique image sequence (consisting of the same set of Mooney and grayscale images). We thus tested our model with the same 12 image sequences presented to human subjects. This resulted in 12 error sequences for humans and model, respectively. We then asked whether there is similarity between these error patterns, as measured by Pearson’s r (Fig. 5d). We found that the model showed a high but imperfect self-agreement at r = 0.78 (p<0.05, Wilcoxon signed-rank test, n=66, pairwise between the 12 model instantiations). Human subjects also showed a significant agreement between each other at r = 0.41 (p<0.05, Wilcoxon signed-rank test, n=66). Importantly, although lower than the level of agreement between human subjects, the model showed significant error pattern similarity to humans at r = 0.29 (p<0.05, Wilcoxon signed-rank test, n=144). This result suggests that our DNN exhibits behavioral alignment to human subjects and recognizes the same images as humans do.

To evaluate whether the internal representations of the model contain information relevant to how human subjects recognize the Mooney images, we used model internal features to predict human subjects’ learning outcomes for individual images (learned vs. not learned). Accurate prediction of human learning outcomes would suggest that the model extracts features that are relevant to humans’ learning success. Using the model’s representation features for grayscale images, prediction accuracy for human subjects’ Mooney image learning outcomes was significant from the 2^nd^ layer onward (Fig. 5e, gray; the 1^st^ layer has index 0), and increased monotonically from early to late layers with a peak AUROC of 66%. The visual features extracted from pre- or post-Mooney images are also significantly predictive of human learning outcome in certain layers, but not as predictive, with post features reaching 59% and pre features reaching 56%. Overall, these results show that the features extracted by the model from the prior-inducing grayscale image are highly predictive of humans’ learning success rate.

Finally, we tested whether the model shows similar invariance properties as human subjects (Fig. 2a-e). To this end, we fed shuffled blocks of image to the model (see Fig. 1a), with random manipulation applied to the grayscale image, and recorded the model’s recognition performance. The model’s performance in the grayscale-manipulated condition was then compared to the original condition or the catch condition, similar to the human experiment. The results are shown in Supplementary Fig. 12. Similar to human subjects, the model exhibits invariance to orientation, size and position manipulations of the grayscale image, as evidenced by a significant interaction effect when comparing each manipulation condition to the catch condition (all ps < 0.001). Because our model was never designed to capture the invariance properties directly, the emergence of invariance in the model’s one-shot perceptual learning ability is nontrivial and adds to the evidence that our model captures the human one-shot perceptual learning phenomenon behaviorally.

### The model suggests that prior-related information is concentrated in HLVC

Armed with a DNN model that recapitulates one-shot perceptual learning ability of humans, has human-aligned error patterns, and predicts human learning success at an image-to-image level, we next used to model to shed light on the computational mechanisms implemented in the human brain. Because the model contains explicit representation of the prior information, we asked which brain region contains neural code similar to the prior information learnt by the model. To this end, we compared the prior information accumulated in our DNN model, which guides the model’s learning behavior, with human brain activity recorded during Mooney image task performance measured by 7T fMRI (N=19; data from Ref ^24^), and assessed the ability of prior information encoded in the model to predict voxel-level neural activity in each brain region.

Given the same sequence of images presented to the human subjects, we predicted each subject’s neural activity using the model’s internal features representing accumulated visual information (state component; see Supplementary Fig. 10 for details), and compared this to a set of baseline predictions. These baselines were obtained from counterfactual catch trials — image sequences that mimic the task format but offer no stimuli for encoded priors, similar to ‘catch’ trials in the psychophysics experiment (Fig. 6a, left; see Methods for details). Since the model’s state component encodes information related to the visual prior information, the improvement in brain prediction score (see Methods for details) as compared to the catch image sequence (shown in orange in Fig. 6a, pooled across all images) indicates the utilization of information related to perceptual priors. We found that fusiform cortex (FC), a region that is part of HLVC, contained the highest proportion of voxels containing prior-related information (29.7%), followed by DMN (13.9%) and FPN (11.2%) (Fig. 6a, right). Outside of FC, we observed a steady increasing trend from early visual regions (<5%) to higher level regions like DMN.

**Figure 6:**
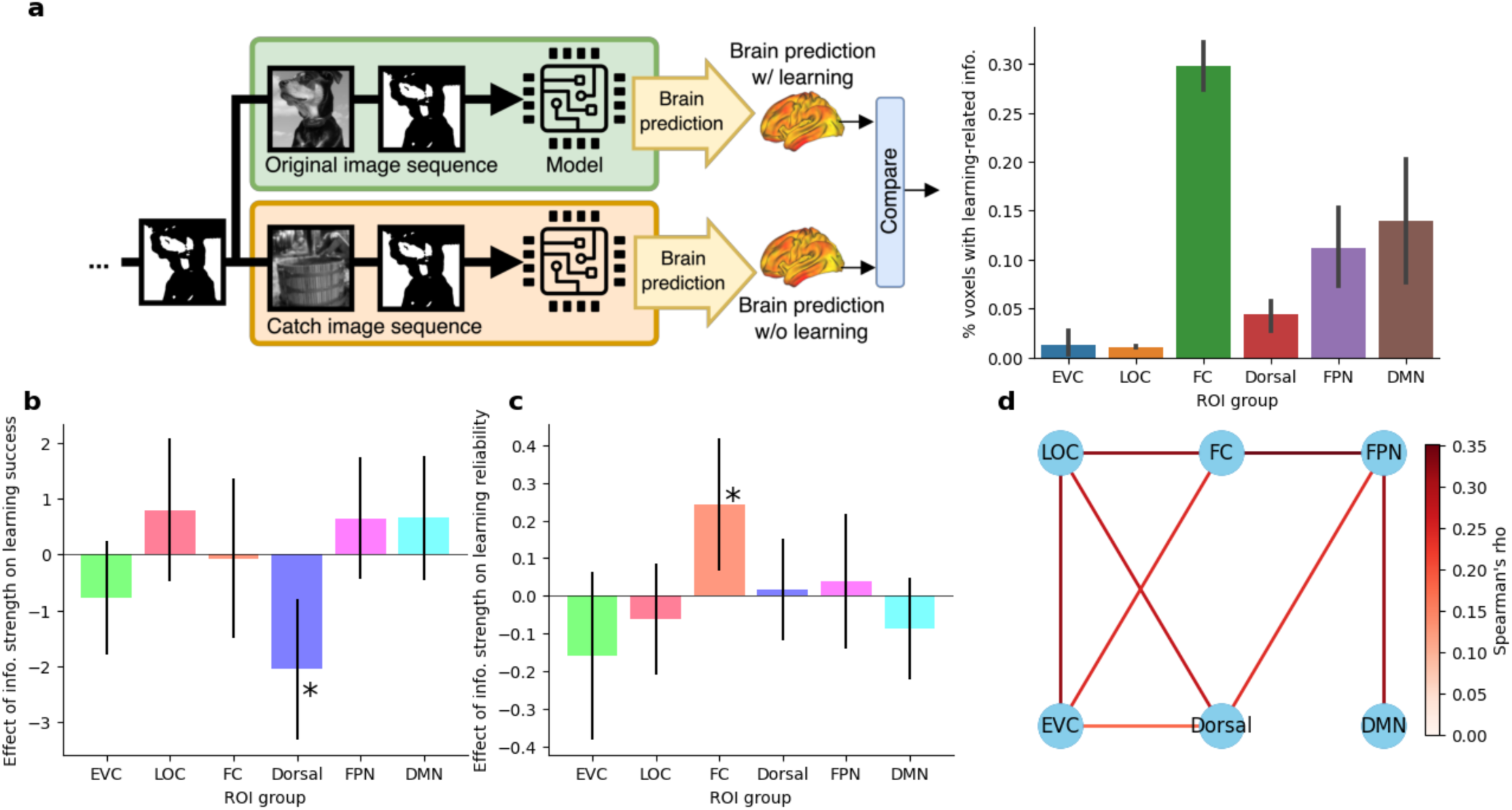
Brain prediction contrast reveals FC’s strong involvement in learning. (**a**) Percentage of voxels in each ROI that show significant improvement over baseline in prediction score at a group level (Pearson’s r, TFCE 10k permutation, p<0.05). Baseline brain prediction is obtained by feeding alternative sequences of images into the model where no learning happens. FC shows highest percentage of significant voxels. (**b**) Higher dorsal stream median information strength is associated with lower learning success. *: significant parameter, error bars: 95% CI of parameter estimate (logit link Binomial family GEE parameter t-test, p < 0.05, FDR-corrected). (**c**) Higher FC median information strength is associated with higher learning reliability. *: significant parameter, error bars: 95% CI of parameter estimate (log link gamma family GEE, parameter t-test, p < 0.05, FDR-corrected). (**d**) Information strength connectivity pattern associated with successful perceptual learning effect (Spearman’s rho parameter estimate using GEE, t-test p < 0.05, FDR-corrected).

### Fusiform cortex information strength assessed by the model predicts the learning effect in humans

Lastly, we evaluated whether successful perceptual learning in humans is related to the strength of learning-related information as measured by the model in each ROI. To measure this information strength, for each image we quantified the proportion of decrease in brain activity prediction error for a typical image sequence compared to the catch image sequence (which induced no learning). We then pooled these results across images at the ROI level. Following earlier work^24^, we defined successful learning in human subjects as 4 or more (out of 6) presentations reported as recognized in the post phase of a Mooney image. We found that the learning-related information strength in the dorsal visual stream is negatively related to the subject’s successful perceptual learning (Fig. 6b), with an increase in dorsal stream information strength from the 50^th^ percentile to 100^th^ percentile reducing average subject perceptual learning success rate from 81% to 61%. This suggests that prior-related information in the dorsal visual stream is inversely related to learning success, a surprising result that hints at a potential competition between the dorsal and ventral visual stream.

We also evaluated whether the reliability of successful perceptual learning in humans is related to the learning-related information strength in an ROI. Taking only the successfully learnt images as defined above, we measured reliability as the proportion of post-phase images that are reported as recognized (varying from 4/6 to 6/6), with 100% being always recognized in the post-phase (hence, most reliable). We found that fusiform cortex (FC)’s information strength was positively associated with the reliability of the perceptual learning effect (Fig. 6c), with an increase in FC information strength from the 50^th^ percentile to 100^th^ percentile increasing the average subject perceptual learning reliability from 84% to 95%. No other ROI’s information strength was associated with success rate or learning reliability, suggesting that the perceptual learning effect is specifically related to information present in FC.

We also evaluated whether the perceptual learning effect is associated with interactions across ROIs by investigating pairwise connectivity between ROIs, where connections are defined by correlation of learning-induced information strength across different images. We found that when an image is successfully learned, there is significant information connectivity across the entire brain network, with fusiform cortex being a central node. Specifically, FC is connected with both EVC and FPN, with FPN further connected to DMN. LOC occupies a more peripheral location in the network graph, being connected only to other visual regions (Fig. 6d, p < 0.05, FDR-corrected). An alternative but weaker path exists from EVC to FPN through the dorsal stream. By contrast, when the image is not learned, we only observed significant connectivity between the dorsal stream and FPN, and no other connections were significant (not shown).

Together, these DNN-informed results demonstrate a central role of fusiform cortex in representing prior visual information and predicting human subjects’ learning success for a specific image.

## DISCUSSION

“Aha moments”, flashes of insight, and other phenomena of one-shot perceptual learning are mysterious and impressive feats of the human brain. Despite decades of research, the site of plasticity and learning underpinning one-shot perceptual learning—fast, long-lasting learning effects in the perceptual domain—remained unknown, in large part due to the learnt prior knowledge being encoded in latent synaptic connectivity (so as to be robust and long-lasting) and difficult to measure using neuroimaging approaches that only capture active neural dynamics.

Here, using convergent approaches from psychophysics, neuroimaging, intracranial recordings, and deep learning, we pinpointed the human HLVC as the seat of neural plasticity subserving one-shot perceptual learning and revealed the potential involved computational mechanisms. The information content of perceptual priors, assayed by psychophysics, uniquely matched the neural coding properties of HLVC, measured by fMRI. Using iEEG, we found that HLVC was the brain region showing the earliest-onset neural signature of prior-guided stimulus processing, suggesting that the latent priors may be encoded and reactivated locally within HLVC. Finally, a vision transformer-based DNN incorporating top-down feedback that shapes visual processing with accumulated prior information was able to recapitulate the one-shot perceptual learning phenomenon in humans and predict the image-to-image human recognition outcome, and the accumulated prior information in the model had the highest correspondence to neural representations in the human HLVC. These multiple strands of converging evidence point to a crucial role of HLVC in one-shot perceptual learning.

Our fMRI experiment was carried out under passive viewing of grayscale images to investigate which brain region has neural coding properties compatible with the invariance properties of perceptual priors uncovered by the behavioral experiment. The logic here is that viewing the grayscale image leaves a ‘trace’ in the activated neural populations, and if the corresponding Mooney image is presented a while later, disambiguation of the post-Mooney image happens, which thereafter becomes a long-lasting memory through consolidation processes. The exact cellular mechanisms involved remain unclear and await future study (we conjecture that the activity trace might be similar to activity-silent working memory^57^). Nonetheless, the invariance properties of neural activation during passive viewing of grayscale images should be equivalent to the invariance properties of priors stored in the one-shot perceptual learning task, because the latter is inherited from viewing of grayscale images during the one-shot perceptual learning task. We note that the same logic was adopted in a long line of research on visual perceptual learning (VPL)— slow, gradual perceptual learning in the visual domain^2^.

Conventional wisdom holds that one-shot learning requires the hippocampus, but a recent lesion study^4^ ruled out this possibility for one-shot *perceptual* learning and instead placed it under the phenomenon of priming^58^. Our observation that one-shot perceptual learning is invariant to size manipulation is reminiscent of earlier studies showing size-invariance in both priming^59,60^ and VPL involving object recognition^30^. Up until now, the relationship between priming, VPL, and one-shot perceptual learning at a mechanistic level has been unclear, with studies on priming focusing on changes in neural activity magnitudes before and after exposure^61^, and studies on VPL focusing on delineating plasticity at different levels of the visual hierarchy^2,28,62^. Our results are compatible with the view that priming and perceptual learning lie on a continuum^28,63^, with one-shot perceptual learning being a special case of priming that has especially long-lasting effects, and a special case of perceptual learning with an especially fast acquisition phase. Interestingly, while three-year old children have similar magnitudes of priming effects as college students, one-shot perceptual learning ability does not reach adult level until adolescence^19,20^. This raises the intriguing possibility that one-shot perceptual learning relies on a perceptual system already fine-tuned by experience.

Previous neuroimaging studies found widespread changes in stimulus-driven neural activity following one-shot perceptual learning^3,22,24–26^ and could not pinpoint where learning takes place in the brain. Using intracranial recordings sampling widespread cortical networks, we observed that neural activity changes driven by prior knowledge emerged first in HLVC, prior to similar changes in EVC, suggesting that top-down feedback from HLVC to EVC could have carried prior-related information. An early primate study using a similar task reported fast changes in IT neuronal firing rates after learning, but did not reveal the time course of these neural activity changes or assess other cortical regions. Interestingly, we did not see a similar neural activity shift towards the relevant prior in higher order brain regions, including FPN and DMN, where such shifts were previously observed in fMRI^24,26^. This is likely due to differences in the recording modalities—high-gamma power is well known to reflect local population neuronal firing rates, whereas fMRI signal can also reflect synaptic inputs and field potential changes uncorrelated to firing rates^64^. Importantly, combining evidence from psychophysics, iEEG and modeling, the present study underscores the key role of HLVC in one-shot perceptual learning, and updates a previous proposal based on fMRI suggesting that the prior knowledge learnt from one-shot perceptual learning is encoded in FPN and DMN.

As part of this investigation, we derived a novel transformer architecture to model the one-shot perceptual learning phenomenon based on top-down mechanisms that convey learnt prior information. We showed that learnt prior information in the model is similar to that contained in the human HLVC, and that the existence of this type of information in the human HLVC predicts more reliable learning in humans. Our network analysis offers a hypothetical mechanism as to how priors shape the perceptual processing. We hypothesize that higher order regions such as FPN might serve as a controlling center for the usage of prior information, which is stored and activated in HLVC and then communicated to other visual areas such as EVC through top-down feedback. The dorsal visual stream, on the other hand, is associated with a lower learning success rate when its information strength is high, suggesting a potential competitive role with the ventral visual stream, consistent with our overall conclusion that the HLVC is critical to one-shot perceptual learning.

Interestingly, while we were developing our model, several similar architectures were described in the machine learning literature that bear a strong resemblance to our model conceptually but were motivated by purely computational considerations with regards to extending the sequence length of transformer-based models (RMT, TransformerFAM, and infini-attention^65^). We see this as a broadly encouraging development in line with other work^66–68^ suggesting a convergence between computational neuroscience research and deep learning.

This work is not without its limitations. First, although our DNN model can predict image-to-image human recognition outcome, its behavioral deviation from human subjects is still appreciable, potentially due to the absence of additional mechanisms (such as the separation between dorsal and ventral pathways) that we do not account for. More accurate understanding of the one-shot perceptual learning phenomenon can inform the development of better computational models that can explain individual human brain activity patterns and learning effects. In addition, in our modeling efforts, we focused on the storage and retrieval of content-specific priors purely based on activation changes, rather than model weight changes. For improved modeling of the long-term retention of learned perceptual priors, model weight updates might be necessary.

Second, the circuit- and cellular-level mechanisms supporting learning-related plasticity in HLVC remain to be uncovered. HLVC is known to support slow, gradual visual perceptual learning (VPL) that occurs at the object level^1,30^. A recent study also revealed that IT neurons encode familiarity—a form of long-term episodic memory^27^. An open question for future investigation is whether the neural code in IT cortex for slow VPL, one-shot perceptual learning, and long-term episodic memory rely on the same or overlapping group of neurons and, if so, whether the neural subspaces representing these distinct types of memories are orthogonal or correlated. In addition, one-shot perceptual learning effects persist for months to years, and, therefore, consolidation of the learnt prior knowledge is likely required and its detailed mechanisms remain to be investigated.

Finally, although we employed a wide range of grayscale image manipulations to delineate the information content encoded in the priors, additional manipulations are possible and can be investigated in future studies. A related question is whether one-shot perceptual learning of low-level visual features, which—although rare—exists in special case scenarios^63^, might rely on other visual regions such as EVC.

## Conclusion

Human perceptual learning is a critical type of learning in humans allowing us to modify how we perceive the world without radically shifting the underlying concepts used to perceive it. One-shot perceptual learning is the crown jewel of this general ability. Our work, localizing the underlying learning process to the HLVC and capturing the learning phenomenon in a DNN with top-down feedback, sheds light on this impressive human feat both biologically and computationally. We anticipate that our work will inspire further research into these novel mechanisms of one-shot learning and support the development of AI models with human-like perceptual mechanisms and computational properties. Furthermore, since altered one-shot perceptual learning reflecting an over-reliance of perception on prior knowledge is observed in multiple neuropsychiatric illnesses involving hallucinations^69,70^, our findings help to pave the knowledge foundation to better understand the pathophysiological processes contributing to these perceptual disorders.

## METHODS

### fMRI – Definition of ROIs

For the ventral and dorsal visual streams (V1-V4, LO1-LO2, IPS0-IPS5, and SPL), ROIs were defined using anatomical masks from a probabilistic atlas that used retinotopy to map ROIs^71^. All overlapping voxels were removed using a winner-take-all approach. Fusiform cortex (FC) was obtained from the “temporal occipital fusiform cortex” partition defined by the Harvard-Oxford atlas. Only voxels that had a >3% probability of belonging in the ROI were included. FPN and DMN ROIs were derived from task-driven activity patterns (GLM and decoding results contrasting pre- and post-Mooney images, respectively) from an independent dataset reported in a previous paper^24^. Specifically, FPN was defined using a binarized statistical map from a whole-brain searchlight decoding analysis of unrecognized pre-Mooney vs. recognized post-Mooney images. DMN was defined using a GLM contrast of the learning effect (unrecognized pre-Mooney vs. recognized post-Mooney) that has been observed in multiple papers^3,24^. Previous control analyses have shown that the results obtained using these ROIs were similar to those obtained using FPN and DMN ROIs from a resting-state atlas^24,26^. For all ROIs, both hemispheres were combined for analysis. The analyses were performed in each subject’s T1 space, with the ROI transformed to this space.

### fMRI – Neural Distance Analysis

Invariant object representation was quantified by the measure of neural distances within the same unique image, across manipulations. t-values from the GLM, aligned to the subject’s T1 space, were used in this analysis^72^, and neural distance metrics were calculated using the rsatoolbox in Python^73^. To increase the signal to noise ratio, randomly chosen pairs of runs were binned into a single run, with t-values averaged across them, creating 8 total “runs” for cross validation. To ensure robustness of the distance metric in RSA, we used cross-validated (c.v.) Euclidean distances as the unbiased distance estimator^73–75^.

From each fMRI run, we construct one Q × P matrix for each ROI (or voxel cluster, in the searchlight analysis), where each of the Q rows is the activity pattern in response to a specific image input and P corresponds to the number of voxels. Here, Q = 80, corresponding to 10 image exemplars × 8 conditions (for details, see SI Methods, fMRI Experiment). Then, we used a leave-one-run-out cross-validation scheme to compute neural distances between images across different folds, as follows:

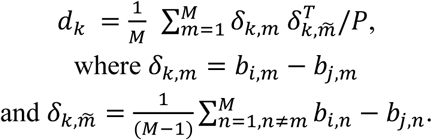

We define a distance metric (d_k_), where m indexes the left-out run, and n indexes the rest of the runs. From the Q × P matrices we have: for the m^th^ run, *b_i,m_* is a 1×P vector for the i^th^ image input and *b* is a 1×P vector for the j^th^ image input. The first term, *δ_k,m_* is the difference in activity patterns between them (*b_i,m_*, *b_j,m_*) in the left-out m^th^ run. The second term, *δ_k,m̃_*, is the averaged difference in activity patterns between the same two image inputs for the rest of the runs. The inner product between *δ_k,m_* and *δ_k,m̃_* are averaged across all M folds, normalized by the number of voxels (P) in each ROI or voxel cluster.

Cross-validated Euclidean distances have been shown to be unbiased distance estimators, such that they are not conflated with noise^74,75^. This is because we can assume noise to be independent across partitions, so any measured noise between *δ_k,m_* and *δ_k,n_* should point in random directions in the high-dimensional space, and therefore be near-orthogonal. Taking the dot product of *δ_k,m_* and *δ_k,n_*, noise should cancel out.

Because this analysis aimed to identify neural invariance properties that matched the invariance properties of the perceptual priors identified in Experiment 1, the line drawing condition was excluded from the analysis, leaving 7 manipulation conditions in total, including the original images (Fig. 3b). Using the 70 image inputs (10 image exemplars × 7 conditions), a 70 × 70 RDM is created using the c.v. Euclidean distances computed for every pair of image inputs (Fig. 3b). Then, to compute within-image, between-condition neural distances (green bars in Supplementary Fig. 5b), the values were averaged across all pairs of manipulation conditions for each image exemplar, then averaged across the 10 image exemplars (green squares in the matrix in Supplementary Fig. 5a). In total, for each subject, 210 values were averaged within each green bar of Fig. 3c, including 21 condition pairs × 10 image exemplars.

### fMRI – Model-based RSA

A model RDM was created to capture the invariance properties of perceptual priors identified in Experiment 1. To this end, we created a 70 × 70 model RDM, in the same layout as the neural RDM (Fig. 3b). Because size manipulation did not significantly impact the learning effect, while orientation and position shift manipulations significantly degraded the learning effect, we assumed the distance between original and size-manipulated conditions to be low (distance = 0, in navy), those between original and rotation/inversion/position shift conditions to be intermediate (distance = 0.5, in teal), and distances between different image exemplars to be high (distance = 1, in yellow). The model RDM was correlated with neural RDMs from each ROI using Kendall’s Tau-B. For statistics, a null distribution was created by shuffling image labels (across both exemplar image and condition labels). For each ROI, the real correlation value was compared to the null distribution to obtain an empirical p value (right-tailed).

A whole-brain searchlight analysis was run in each subject’s T1 space, using a 6-voxel radius sphere size. For each voxel cluster, the neural RDM was correlated with the model RDM using Kendall’s Tau-B. The correlation values were transformed into Fisher’s z, and normalized into standard MNI space. For group-level analysis, the z-value maps were spatially smoothed (12 mm FWHM) and submitted to a one-sample t-test across subjects. Significance was assessed by permutation (using FSL randomize) and threshold-free cluster enhancement (TFCE) method^76^, thresholded at a p < 0.05, FWE-corrected level.

### iEEG Data Analyses

Electrodes were assigned to an ROI if their MNI coordinates lay within a voxel in that ROI. FPN and DMN ROIs were defined as above, from ^24^. EVC, HLVC, and Dorsal ROIs were defined as above, except no voxels were excluded for low probability, and there is no prohibition that a voxel be in only one ROI. The Limbic ROI was defined as the union of the “Cingulate Gyrus, anterior division”, the “Frontal Orbital Cortex”, and the “Insular Cortex” regions of the Harvard Oxford Anatomical Atlas.

The electrode mean activation time course for each condition (Pre, Post, Grayscale) was calculated as the mean across trials for that condition and electrode. The mean time courses for each ROI (Fig. 4, top) were calculated as the average across electrodes within the ROI, for each condition. Error clouds were calculated as the paired SEM^52,53^ across electrodes at each timepoint. Significance testing was performed using a paired t-test at each timepoint (one-tailed, to identify time points with higher HGP in the post than pre stage). We focused on the post > pre effect here to exclude potential task difficulty-related effects, since the pre phase is more difficult, whereas the post phase has heightened prior- and recognition-related processing. To correct for multiple comparisons across time points, a cluster-based permutation test was used (see below). Since trials terminated at image offset when a response was given, the number of trials with data varies across timepoints, especially towards the end of the trial. Once all trials of one condition for an electrode have terminated, the electrode is dropped from that and subsequent timepoints in corresponding mean ROI time courses. See Supplementary Fig. 7b for each ROIs’ electrode survival time course.

The Image Preference Analysis quantifies how well each electrode’s image selectivity during Pre and Post conditions aligns with its image selectivity during Grayscale image trials (Fig. 4, bottom). For each electrode, the mean HGP time course for each image was calculated, separately for the grayscale, pre, and post phases. At each timepoint, the relative values of each image’s mean HGP in the mean Grayscale time courses defined an image sorting order. The Pre (/Post) images’ mean HGP values at that timepoint were arranged in the same image sorting order and a best fit linear regression line was calculated. The best fit line’s slope was extracted, and this procedure was repeated for every timepoint, producing a time course of slopes for the Pre (/Post) condition at each electrode. Mean traces, paired SEM, and significance were calculated similarly to that described above, except that Wilcoxon signed-ranked test was used in place of paired t-test. Timepoints at which there are not at least two images with trial data in both the grey condition and the Pre (/Post) condition were dropped. See Supplementary Fig. 7c for each ROIs’ electrode survival time course.

We used an adapted cluster-based permutation test^77^ to accommodate the varying degrees of freedom (DOF) at different timepoints due to the varying trial durations. We defined a cluster’s DOF to be the mean DOF for the timepoints it spans, rounded to the nearest integer. We calculated cluster summary statistics on our real data as the sum of the test statistic for each cluster, and also note the DOF for each cluster. We then produced a permutation-derived null distribution for each DOF using the following procedure. These null distributions were defined so that all data points within one distribution derived from clusters with the same DOF. For each permutation, we identified the largest cluster summary statistic and its DOF, and its summary statistic was assigned to the corresponding null distribution. A count of permutations that produce no significant clusters was maintained and a corresponding number of zero-valued data points was added to each null distribution proportionately to the number of data points it has once permutation generation has ended. Permutations were generated and this process was repeated until there are 1000 data points in all null distributions with DOFs matching the clusters from the real data, or all possible permutations had been produced, or 1,000,000 permutations had been produced. P-values were then assigned to each cluster by calculating the percentage of data points in the null distribution with the corresponding DOF that were greater than the cluster’s summary statistic. This corresponded to a one-tailed cluster-corrected test.

Since massive univariate analysis on individual timepoints (followed by cluster-based permutation test to correct for multiple comparisons) does not statistically assess onset timing, we performed a bootstrap analysis to compare the relevant timings between ROIs in key conditions of interest^78^. For each condition of interest, we generated 2,000 bootstrapped replications of the corresponding analysis described above, yielding 2,000 sets of significant clusters. An estimate of the onset time distribution for each real cluster in a condition of interest was generated by extracting from each replication the earliest significant timepoint that was part of a cluster that overlapped with the real cluster. Only replications with significant clusters that overlapped with the real cluster contributed points to these distributions, and a minimum of 800 points contributed to the estimated onset time distribution for all conditions of interest^79^. The points at the 2.5^th^ and 97.5^th^ percentiles were reported as the 95% confidence interval (CI) onset.

### DNN Model Architecture

Our model consists of two learnable components, a vision transformer and cross-attention module, and a memory state module that is updated via a fixed rule. The vision transformer is initialized from the base sized DinoV2 vision transformer, which has 768 hidden units and 12 layers of self-attention mechanisms for processing visual features. The original vision transformer was pretrained using color images, whereas our task used grayscale images. We handled this difference by duplicating the grayscale intensity to the R, G, B channels so as to not bring additional information into the model. This initial vision transformer was then trained on the image recognition task sampled from the ImageNet1k dataset. To keep training time reasonable, we use low-rank adaptation (LoRA)^80^ to finetune the key and value weight matrices of the self-attention layers only. LoRA learns additional weights that are added to the existing weights, which ensures the visual features already learned is not lost. We use a matrix rank of r=64, and alpha value of 64. The cross-attention module is used to retrieve information from the state module and is fully trainable.

The model operates in two stages, first a feedforward run to produce retrieval queries, and then a conditioned run to generate an object classification given past knowledge using this query. When initially shown an image, the image is fed into the vision transformer without any previous information. This produces a priming feedforward output that encodes visual representations in the input image. The output of size (1+256 tokens, 768 hidden dimensions) is down-sampled to (1+9 tokens, 768 hidden dimensions) using adaptive down-sampling to reduce computation cost and generate a query.

This query is then used to compute the feedback conditioning for the conditioned feedforward run. The previous state (1+10+256 tokens, 768 hidden dimensions) is used as the key and value for the cross attention, while the query generated from the feedforward run is used as the query. This produces a feedback conditioning (1+9 tokens, 768 hidden dimensions) representation as the top-down signaling for the next stage of computation.

The second conditioned feedforward pass then occurs with the feedback conditioning providing conditional information from the prior state to the same vision transformer from the prior stage to generate an output. The conditioning tokens (10) are concatenated with the image patch tokens. The second run produces top-down informed outputs (1+10+256 tokens). The first token corresponds to the CLS token from the model; 10 tokens are from the conditioning tokens; and the 256 tokens correspond to image patches produced from the current input image. Lastly, the state is updated with a moving average with a learned constant during training and kept unchanged during evaluation.

### Notable Differences from Existing Models

There are a few architectural details that can explain the difference in performance. Existing models such as BLT and CORnet rely on convolutional neural networks (CNN)-based layers to extract visual features, whereas our model uses a vision transformer, which encourages more complex interaction between parts of the visual input.

A second crucial difference between our model and BLT and CORnet is how the recurrence is implemented. In our model, the state is a separate module that carries information about the past and gets updated with a simple moving average mechanism. This protects the priors from being overly affected by the current step of activations. On the other hand, BLT and CORnet do not keep a separate state module; all the prior information needs to be stored in the current activation, which sacrifices the expressivity of current visual representation and limits the storage capacity.

Another difference from existing models is that our model is explicitly selective about how prior knowledge affects the current visual processing. Our model achieves this selective retrieval by first computing bottom-up visual features without the conditioning, followed by retrieval of relevant priors using cross attention. Existing models simply merges the past state with the current activation, where prior activations implicitly modulate current activations and lacks fine-grained selectivity of past prior information.

### DNN Model Training and Evaluation

We train the network on a computational adaptation of the Mooney Task. To construct a sequence of image, we first obtain all unique color images from the ImageNet dataset at random, convert them to grayscale images, and apply random thresholding between pixel intensity 50 and 205 to each image to produce the binarized version of them (‘Mooney image’). For each batch during training, we first obtain 3 pairs of grayscale and Mooney images. To encourage the reuse of information across time and not discard information right after usage, we repeat each constructed sequence 3 times. Finally, we shuffle the entire sequence before feeding into the model. This same image sequence structure is used in the evaluation shown in Supplementary Fig. 11a. Below we give an example of a typical sequence used for training the model. We first obtain 3 grayscale images, and notate them as G1, G2, G3. We then binarize them with random threshold and obtain M1, M2, M3. We construct a sequence wherein each M-G image pair occurs 3 times: [M1, G1, M1, G1, M1, G1, M2, G2, M2, G2, M2, G2, M3, G3, M3, G3, M3, G3]

Finally, we shuffle this sequence to encourage the model to be able to handle arbitrary sequence order. E.g., a valid shuffled sequence can be: [G2, G2, M3, G1, M2, G3, M1, G3, G1, M2, G2, M1, M2, M3, M1, G1, G3, M3].

For training, we accumulate gradients for 16 steps with a batch size of 32 sequences on 8× A100 GPUs with the PyTorch lightning framework. All images in the sequence are weighted equally in the cost function. Supplementary Fig. 11a shows model performance during the evaluation phase, using held-out image sequences following the same shuffled structure. All image sequences used during training and evaluation phase had a length of 18. Any Mooney image presented before (/after) the first corresponding Grayscale image is designated ‘Pre’ (/’Post’) in the results plot. Baseline models were trained and evaluated using the same approach.

We next evaluated our model on longer synthesized image sequences (sequence length: 630, using 210 unique Mooney images), again created from grayscale images that were randomly selected from ImageNet, and thresholded using a random threshold between 50 and 205 to generate the corresponding Mooney images. Repetition effect was assessed by presenting two repetitions of the same 210 Mooney image in the same sequence, without the corresponding grayscale images (sequence length: 420). Sequences followed the same block-shuffled structure as the human psychophysics experiment (Fig. 1a). The results from this evaluation are shown in Fig. 5b.

### Model Behavior Comparison with Humans

From an online study, behavioral data was obtained from 12 subjects performing the Mooney Task (see SI Methods). The same 219 Mooney images and their matching grayscale images were presented to each subject in a unique order. We supplied each image sequence used in the human experiment to the model and obtain 12 sets of model behavioral data. When comparing model behavior to human behavior, we first take the top 90 Mooney images on which human subjects showed the strongest perceptual learning effect, measured by the average pre-to-post accuracy increase (the same 90 images were used in the in-person behavioral experiments). We then exclude any images that are not in our model’s label space due to the model’s inability to predict images outside of the label space. This resulted in 78 unique Mooney images (and their corresponding grayscale images) used in the model-human behavior comparison shown in Fig. 5c-d. Equivalent results for baseline models are shown in Supplementary Fig. 11b.

To obtain the results in Fig. 5d, we first transformed each subject’s behavioral output or model output into a binary 1×234 vector denoting correct/incorrect identification of a specific image shown in the pre/gray/post phase. We then calculated pairwise correlations between human subjects and between the 12 sets of model outputs, yielding 66 error pattern similarity measures for each. We further obtained 144 (12×12) model-human error pattern similarity measures.

### Learning Outcome Prediction

Using the same behavioral data set mentioned above, for each subject (N = 12), we removed images that were already recognized in the pre phase. We then obtained the features from the DNN components including the state, query, and the conditioning, and used an SVM classifier with a learnable linear kernel from the sklearn python package to predict whether the subject will recognize the image in the post phase. To avoid data leakage, we applied a 12-fold cross-validation and report the mean test performance over folds. AUROC was used to measure the performance.

### Neural (fMRI) Data Prediction from DNN

To predict the post phase brain beta values (output from the GLM at individual image level, see ^24^), we use a kernel ridge regression model with sparsity alpha=1, using the latent visual portion of the recurrent model state as features (see Supplementary Fig. 10). To obtain a set of beta map predictions without overfitting, we use leave-one-image-out cross validation for each subject. All predictions were made after registering subject-level beta maps to the standard MNI152 space, and predictions from 4 different model seeds were averaged together.

The baseline predictions are obtained from constructing counterfactual model representations, using catch images. The counterfactual representations for an image I is obtained by replacing only the corresponding grayscale image for I (i.e., replacing the correct learning material). The average counterfactual representation over all possible alternative learning material (32 alternative images) is used as the feature.

We first find the correlation between predicted betas and the true betas. We then apply z-transform to the score. Repeating this process for the normal beta predictions and the baseline beta predictions gives us two maps per subject. Subtracting the baseline score map from the normal score map, we obtain the map of score improvement.

To calculate the information strength of a region, we used the following procedure.

Let’s denote the beta prediction for voxel j and image i as *ŷ_ij_*, and the true beta there *y_ij_*. We first calculate the relative error 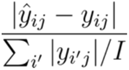 for both the normal prediction and the baseline predictions. This provides us one normalized error map per combination of subject and image. Then we compare the error of baseline and normal predictions. This is obtained with 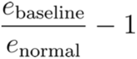. This yields one map per subject and image, with 0 indicating no difference between both performance and higher value indicating improvement in performance. In the following experiments, we only consider maps with strictly non-negative entries in this map. For the ROI level analyses in Fig. 6b, c, d, we find the median information strength within the ROI. The ROIs were as described earlier; here we threshold the ROI masks from the probabilistic atlases at 25% and assigned a voxel to the most likely ROI.

To compare prediction score with noise ceilings, after we obtain the prediction score, we average the z-transformed scores between 4 different model seeds. We obtain lower and upper noise ceiling maps for each subject with the Representation Similarity Analysis (RSA) method^81^. Specifically, to calculate the upper noise ceiling estimate for a voxel center, we first find its average RDM across subjects. The average similarity of this average RDM to all subject is the upper noise ceiling. To calculate the lower noise ceiling, we use leave-one-subject-out similarity. For each subject, we leave their RDM out, take the average RDM across the remaining subjects, and calculate the similarity between the held-out RDM and the average RDM. To compare the score against either lower or upper noise ceiling, we use a permutation-based 2-sample t-test in a whole-brain voxel-wise analysis, with threshold-free cluster enhancement (TFCE) correction with 10,000 permutations. A one-sided comparison with the max statistic was used to produce voxel-level significance values. The results were summarized by ROIs.

To compare prediction score with baseline score, we first average the z-transformed normal prediction scores between 4 different model seeds. We use the same procedure to obtain scores for the baseline predictions. Then we use a permutation based 2-sample t-test with TFCE correction with 10,000 permutations. A one-sided comparison with the max statistic is used to produce voxel level significance values.

To obtain the estimates for the analysis in Fig. 6b, we first obtain a binary label of ‘learned’ versus ‘not learned’ based on the subjective report of the subjects. We fit a binomial family generalized estimating equations (GEE) model grouped by subjects with the logit link function to predict whether the image is learned from information strength in each ROI. The model can be expressed in the R-style formula: learning ∼ FC + EVC + LOC + DMN + Dorsal. After fitting the model, the parameters corresponding to each ROI’s contribution to learning is tested using t-test.

To obtain the estimates for the analysis in Fig. 6c, like in Fig. 6b, we first obtain the reliability score by excluding the images that are not learned, followed by finding the number of post recognitions over the number of occurrences in post phase. We fit a gamma family GEE model grouped by subjects with log link function to predict the reliability, which is between 0 and 1. The model can be expressed in the R-style formula: reliability ∼ FC + EVC + LOC + DMN + Dorsal. After fitting the model, the parameters corresponding to each ROI’s contribution to learning is tested using t-test.

To obtain the estimates for the information connectivity between ROIs, we first calculate spearman’s rho between the pairs of ROIs’ information strength for each image. This produces a connectivity matrix between ROIs for each subject. To aggregate this connectivity across subjects, we take the lower triangle of this connectivity matrix and fit a GEE model grouped by subjects. This model can be written in the R-style formula as: connectivity ∼ C(ROI pair). After fitting the model, the parameters corresponding to each unique pair of ROIs are tested using t-test.

## Acknowledgments

This work was supported by a W. M. Keck Foundation medical research grant (to EKO and BJH), an NSF grant (BCS-1926780, to BJH and OD), as well as NYU Grossman School of Medicine. We would like to thank Kendrick Kay for providing the pRF data analyzed herein, Daniel Haşegan for support with ECoG analysis, and Larry Squire for discussions on the psychophysics experiment. We would also like to thank Michael Costantino and the NYU Langone HPC team for their support of our AI research efforts.

## Supplementary Result

### pRF Analysis

To investigate image manipulations’ potential impacts on neural coding at each stage of visual processing, we employed previously published population receptive field (pRF) data from the human ventral visual stream^1, 2^. A voxel’s pRF measures the center and size of its receptive field based on measured fMRI BOLD signal, reflecting an average property across neurons sampled by that voxel. For each ventral stream region of interest (ROI), we quantified how often a voxel’s pRF contains the diagnostic image feature for both the manipulated and the original grayscale image. Specifically, for each grayscale image and each ventral stream ROI, we first identified all the pRFs that overlapped with the diagnostic image feature (e.g., the dog’s head) in the original image, and then identified the subset of these pRFs that still contained a significant portion of the diagnostic feature in the manipulated image (for details, see Supplementary Methods). The number of such pRFs, as a fraction of the total number of measured pRFs, was calculated for each ROI and manipulation condition.

This analysis showed that in anterior HLVC (mFus), size and orientation manipulations have relatively small impact on neural coding (70–100% pRFs retain diagnostic feature), while VF shifts had relatively large impacts (20–40% pRFs) (Supplementary Fig. 1). In addition, as expected, all manipulations have larger impact on neural encoding in earlier visual regions.

## Supplementary Figures

**Supplementary Fig. 1:**
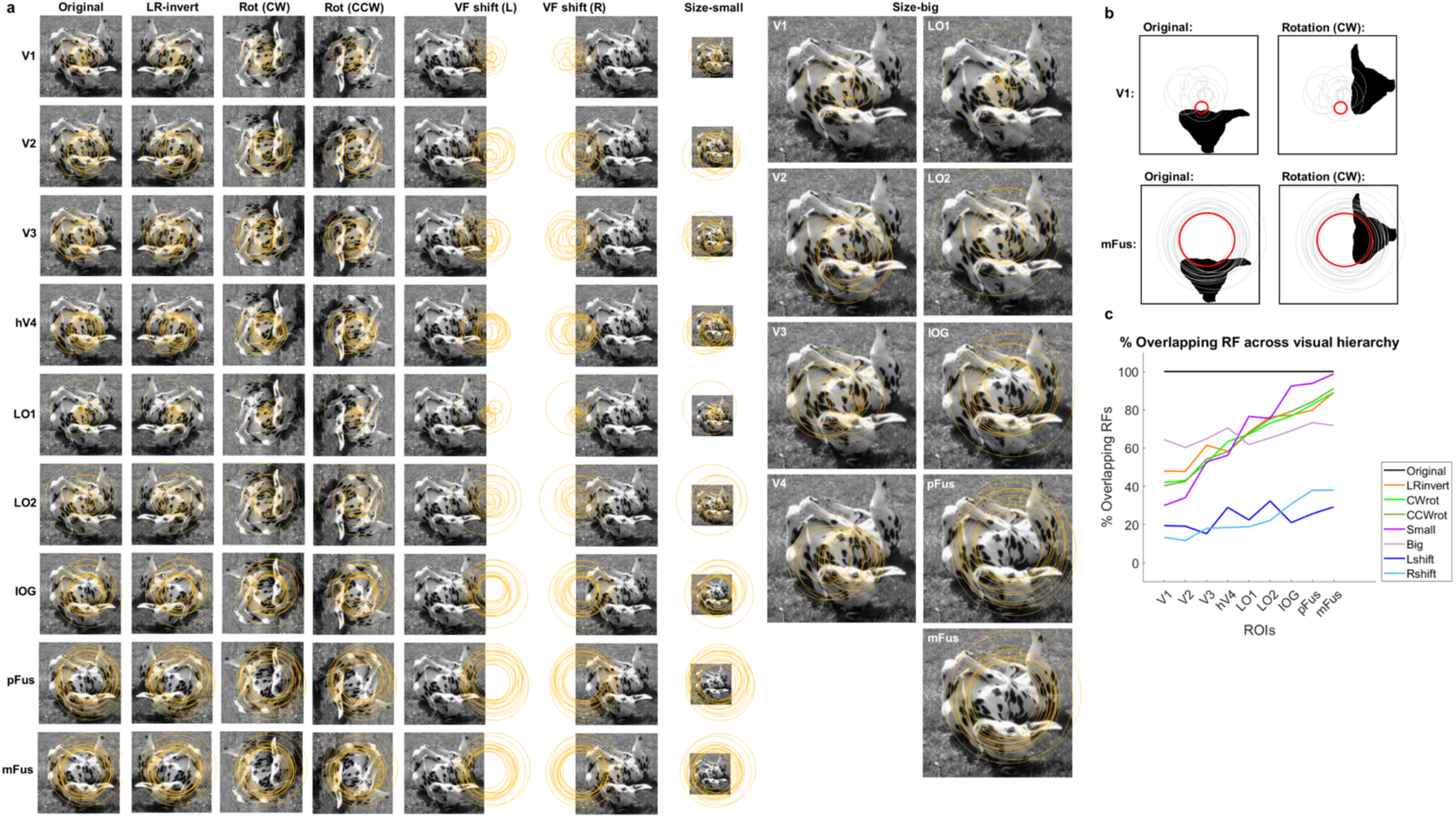
pRF analysis in the context of our psychophysical task manipulations. (**a**) A sample image is shown in all manipulations, overlaid with a randomly selected sample (n = 15) of voxel-level pRFs from each ROI. pRF data shared by Kendrick Kay. V1-V4, LO1, LO2 are from Kay et. al 2013^1^ and defined by a retinotopic localizer; IOG, pFus, and mFus are from Kay et. al 2015^2^ and defined by a functional localizer in conjunction with anatomical landmarks. (**b**) Analysis schematic. Black silhouette indicates key object feature within the image. Black circles are a subset of pRFs for illustration. Red circle indicates a specific pRF from a single voxel being tested for overlap with object feature. This was repeated for all pRFs in each region. (**c**) pRF analysis results. Lines indicate percentage of measured pRFs that retain significant overlap with the diagnostic image feature after each type of image manipulation, using all measured pRFs that have a r^2 value of >50%. In **a** and **c**, ROIs are arranged roughly according to the visual hierarchy, from early visual cortex (V1-hV4) to posterior HLVC, to anterior HLVC.

**Supplementary Fig. 2:**
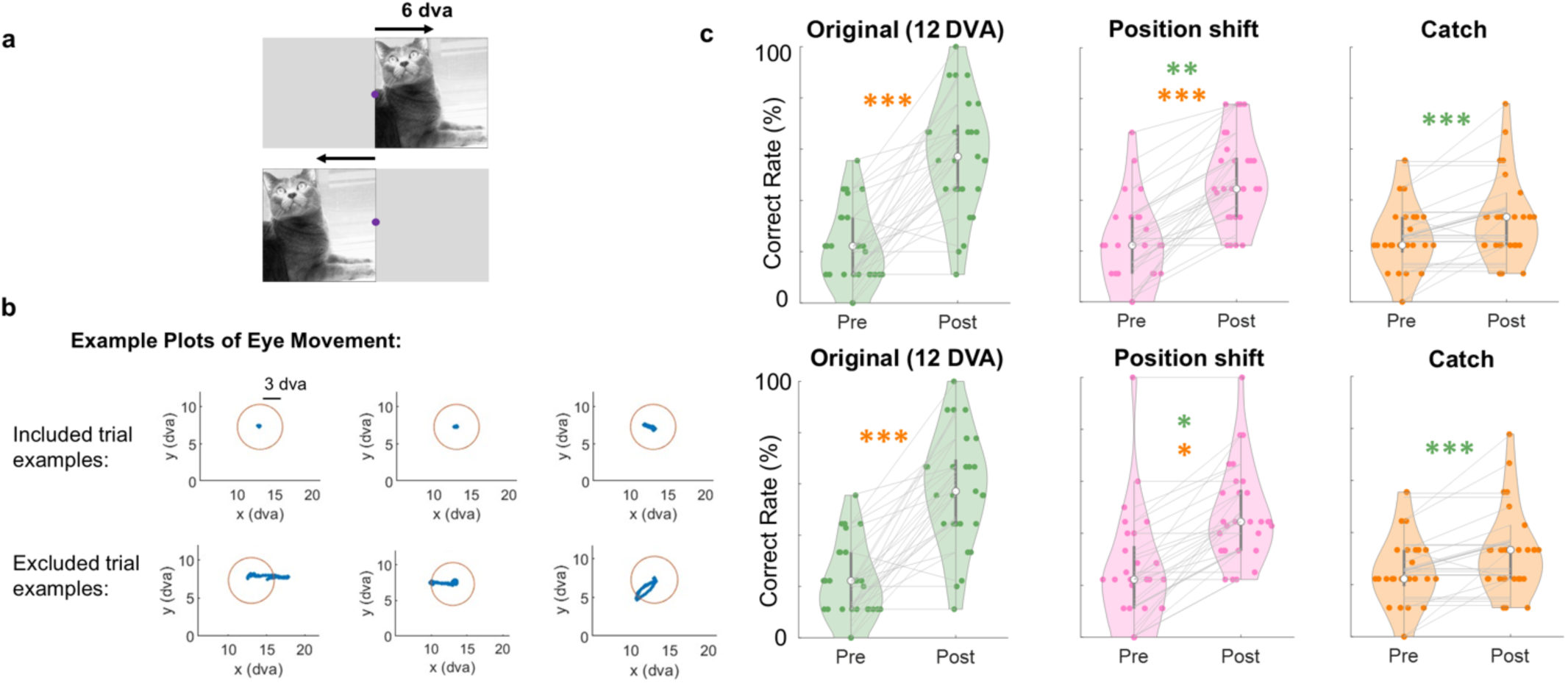
Position shift trials controlled for gaze shifts. (**a**) Image manipulation in position shift trials, shown for an example grayscale image. (**b**) Example plots of eye movements in included trials (top) and excluded trials (bottom). Orange circle indicates the exclusion threshold of a 3 dva radius around central fixation. (**c**) Top: data from all trials (reproduced from Fig. 2a & e). Bottom: only trials where visual fixation was maintained within the 3 dva circle were included. For results in the bottom panels, a generalized mixed effects model was used to account for unbalanced trial numbers, where the dependent variable is the recognition rate; fixed effect is manipulation type, and random effects are subjects and trials. Two subjects did not have eye-tracking data, so all trials were included.

**Supplementary Fig. 3:**
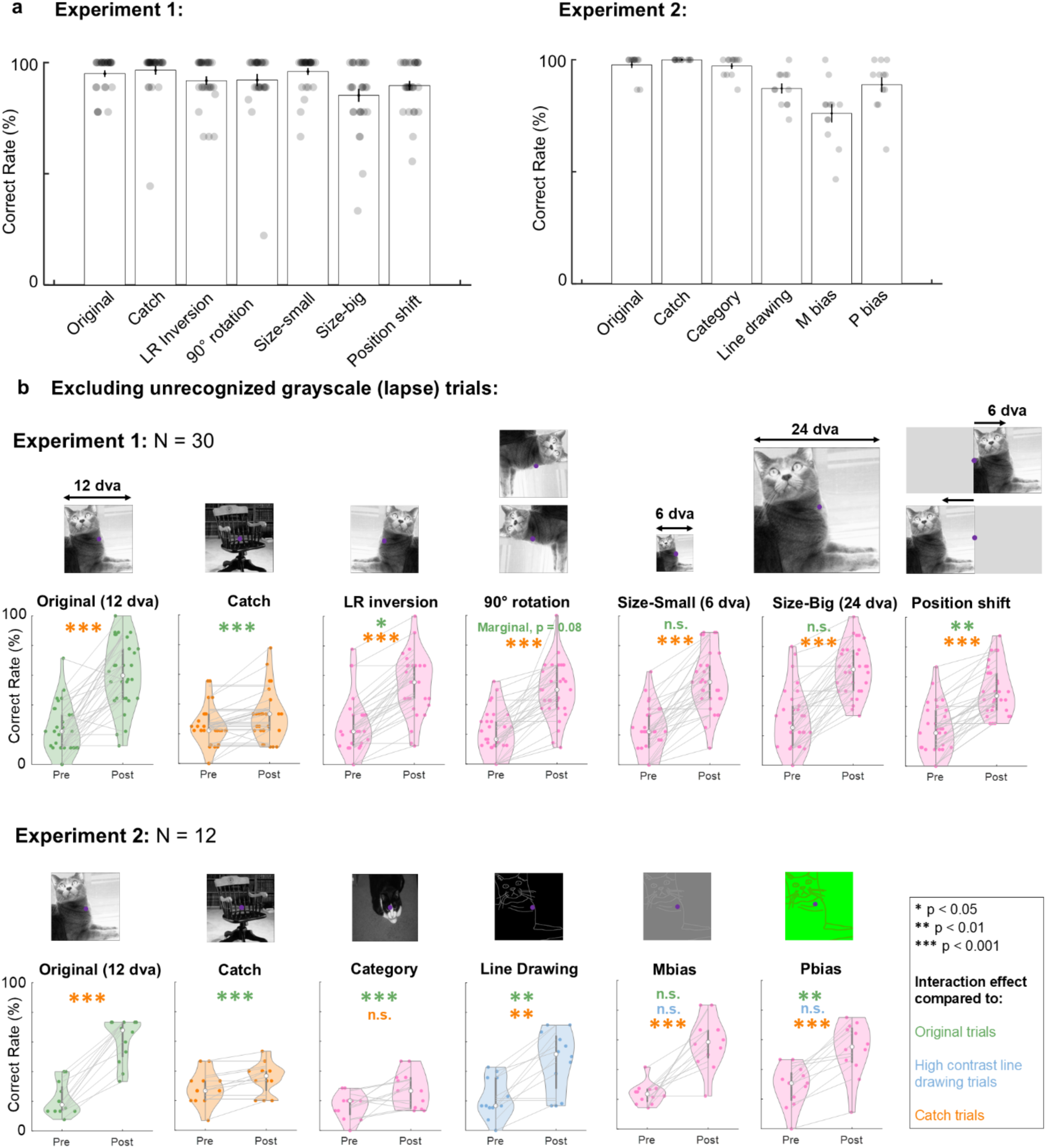
Control analysis including only trials where the prior-inducing image was correctly identified. (**a**) Grayscale image identification rates for each for each manipulation condition in Experiment 1 (left) and Experiment 2 (right). Error bars indicate SEM across subjects. (**b**) Behavior results excluding trials with unrecognized grayscale images. Asterisks denote statistically significant interaction effects compared to original (dark blue), high-contrast line drawing (light blue) or catch (orange) trials. Figure layout is the same as in Fig. 2. This control analysis yielded the same pattern of results for all experimental conditions in both experiments.

**Supplementary Fig. 4.**
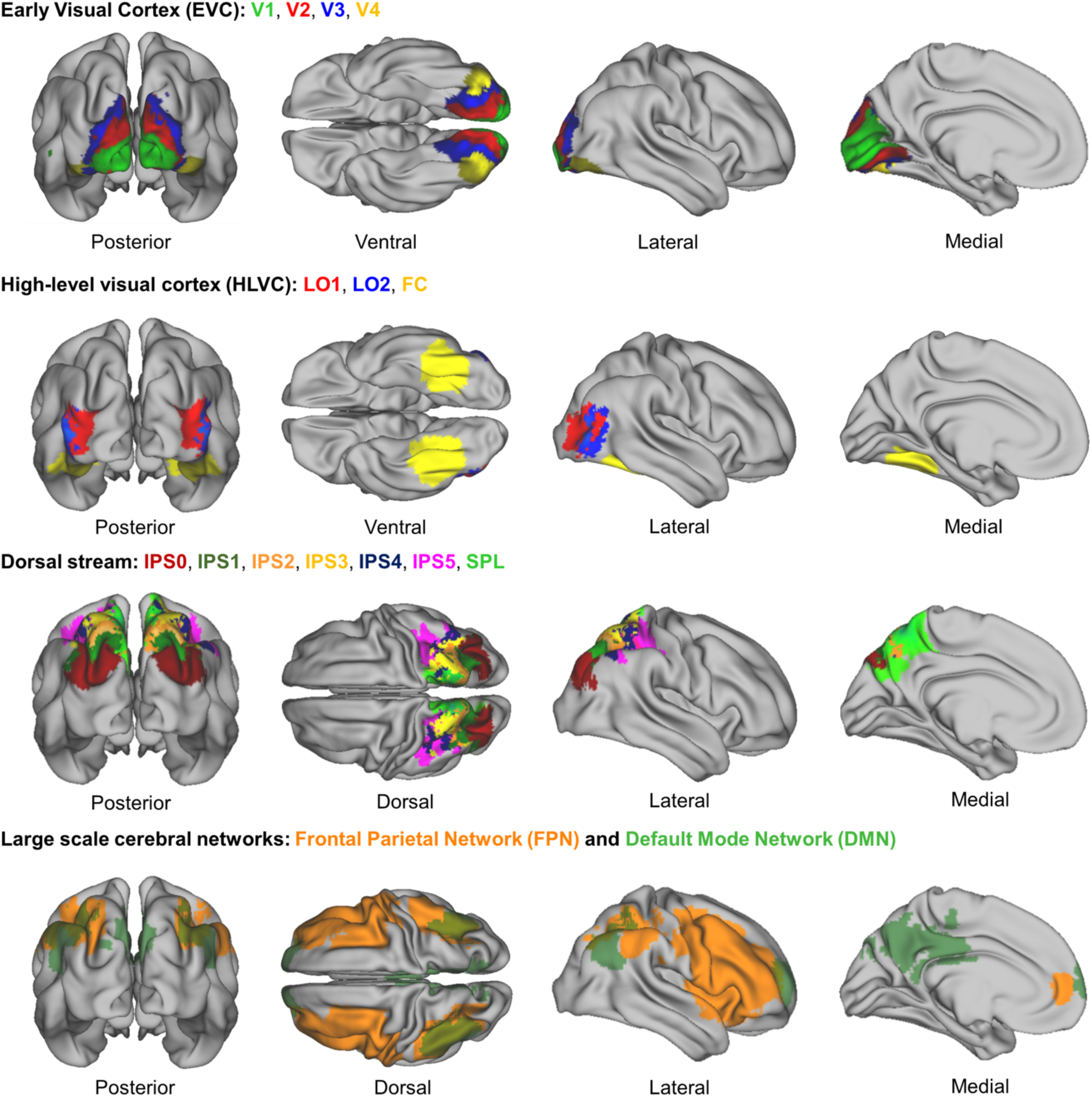
ROIs used in fMRI analysis. V1-V4; LO1 and LO2; IPS0-5; SPL are taken from a probabilistic atlas defined in Wang, et. al (2015)^3^. FC is taken from the Harvard-Oxford atlas (“temporal occipital fusiform cortex”). Frontoparietal and Default Mode Network ROIs are task-defined functional ROIs, taken from Gonzalez-Garcia (2018)^4^.

**Supplementary Fig. 5.**
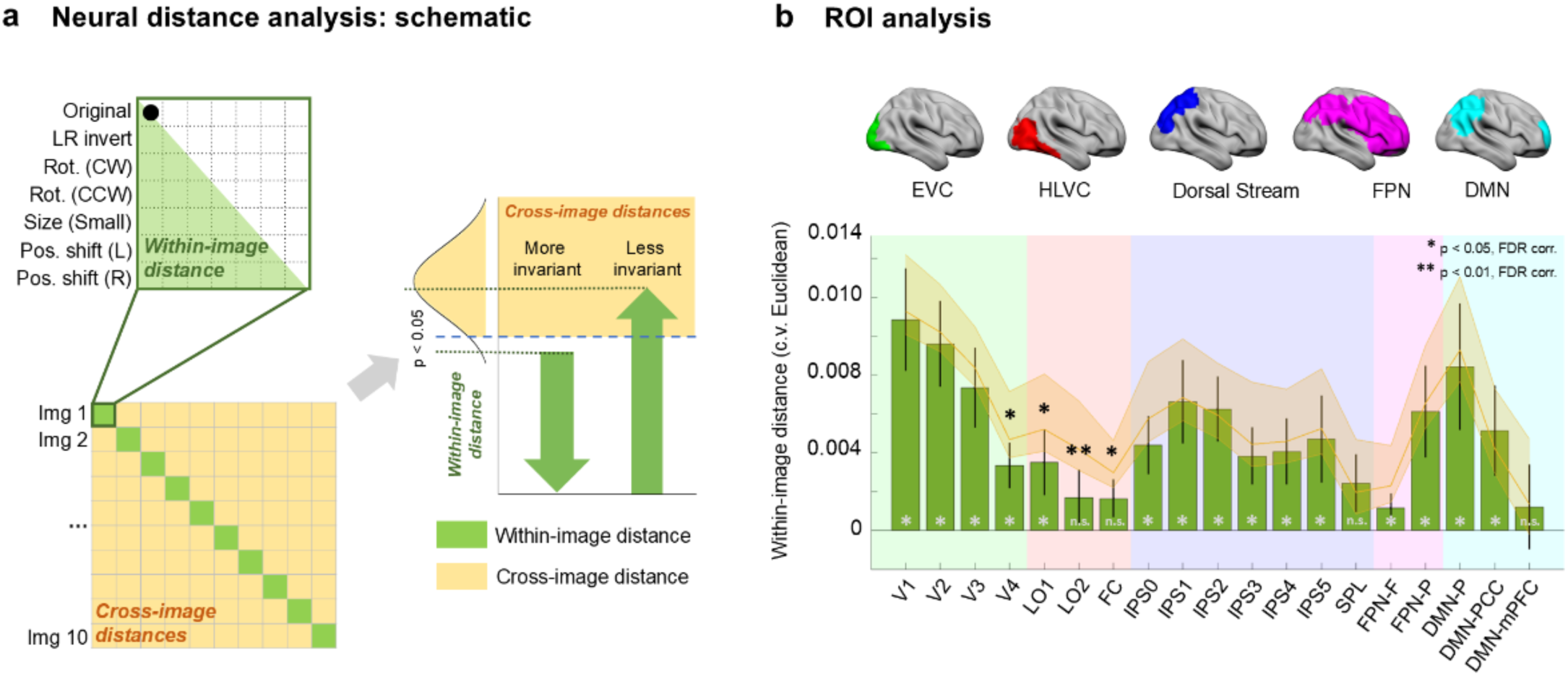
Neural invariance analysis. (**a**) Analysis schematic. The layout of the fMRI RDM is the same as in Fig. 3b. (Left) Voxel-wise fMRI activation patterns are extracted from each ROI, and cross-validated (c.v.) Euclidean distance was calculated across all pairs of image-condition combination to generate a 70 × 70 RDM. (Right) Within-image, between-condition distances were compared to the between-image distances to assess for manipulate-invariant neural representations. (**b**) Top: locations of ROIs, grouped by networks; for detailed ROI locations, see Supplementary Fig. 4. Bottom: within-image, between-condition distances for all ROIs (green bars) and the “noise ceiling” estimated by multiple iterations of subsampling between-image distances with similar condition-pair distributions (yellow). Black asterisks denote significant differences between within-image between-condition distances and between-image distances (FDR corrected). Light gray asterisks denote within-image between-condition distances that are significantly greater than 0 (one-sampled t-test, FDR corrected), where 0 indicates full invariance. Error bars indicate SEM across subjects. See SI Methods section “fMRI Analysis—Neural invariance analysis” for details.

**Supplementary Fig. 6.**
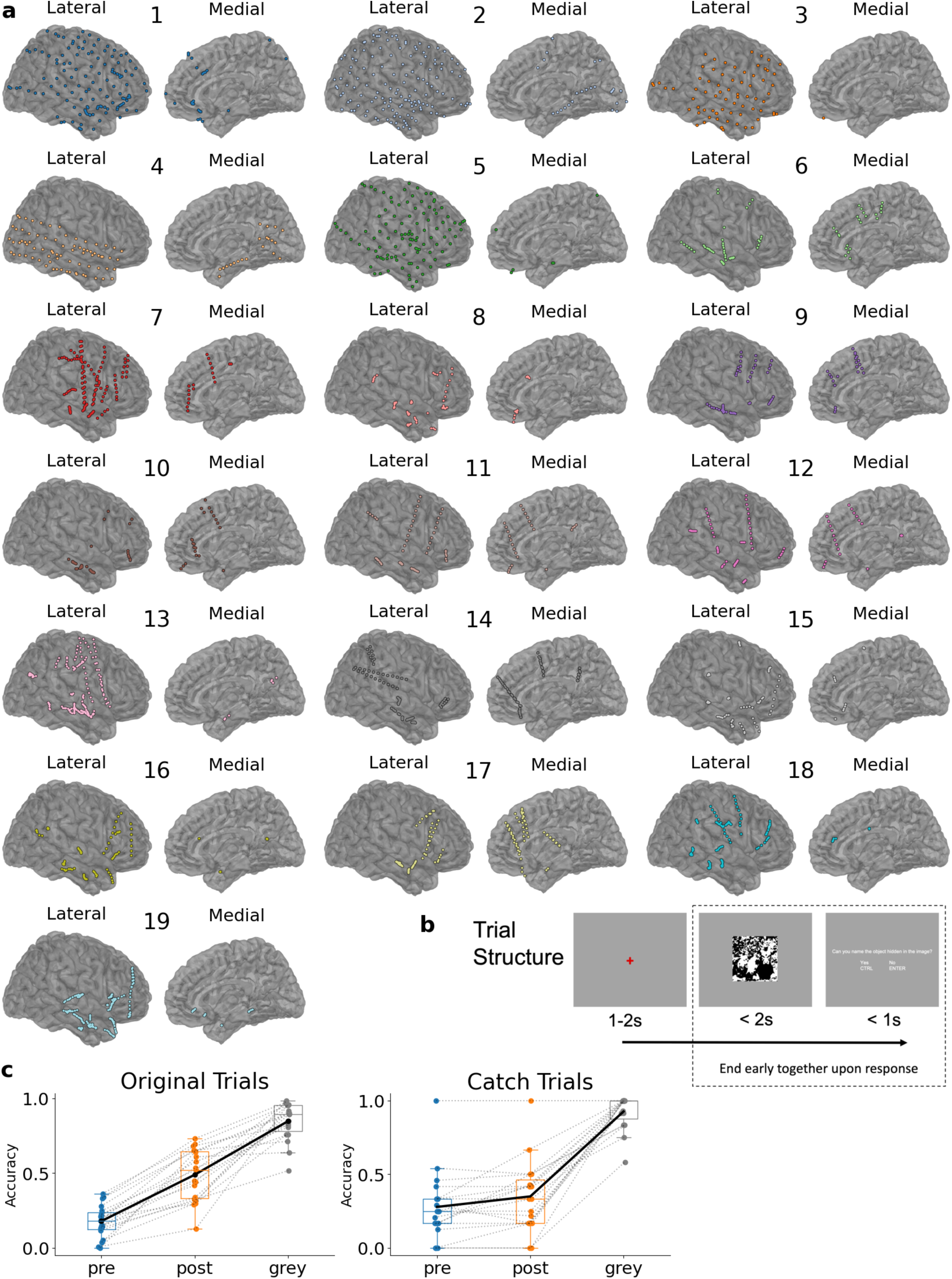
Electrode coverage, trial timeline and behavioral results from iEEG patients. **(a)** Electrode placement for all patients included in the study. Left hemisphere electrodes shown reflected across the midline, such that they are combined with right hemisphere electrodes in the plots. **(b)** Trial timeline for the iEEG task. See Methods section iEEG Experimental Paradigm for details. **(c)** Patient accuracy in recognizing Mooney and grayscale images in the original (left) and catch (right) trials. Box-plot elements defined by the center line at the median, box boundaries at the first and third quartile, and the whiskers at the farthest data point not more than 1.5 times the length of the box away from the box boundaries. Assessed by a paired Student’s t-test, difference in the accuracies for pre and post images is highly significant in original trials (p = 3.3 × 10^−8^); and significant in catch trials (p = 0.026, reflecting repetition-induced baseline effect). A two-way repeated-measures ANOVA ([pre vs. post] × [original vs. catch]) showed a significant main effect for Pre vs. Post (F_1,18_ = 62.0, p = 3.1 × 10^−7^) and a significant interaction effect (F_1,18_ = 34.2, p = 1.5 × 10^-5^).

**Supplementary Fig. 7.**
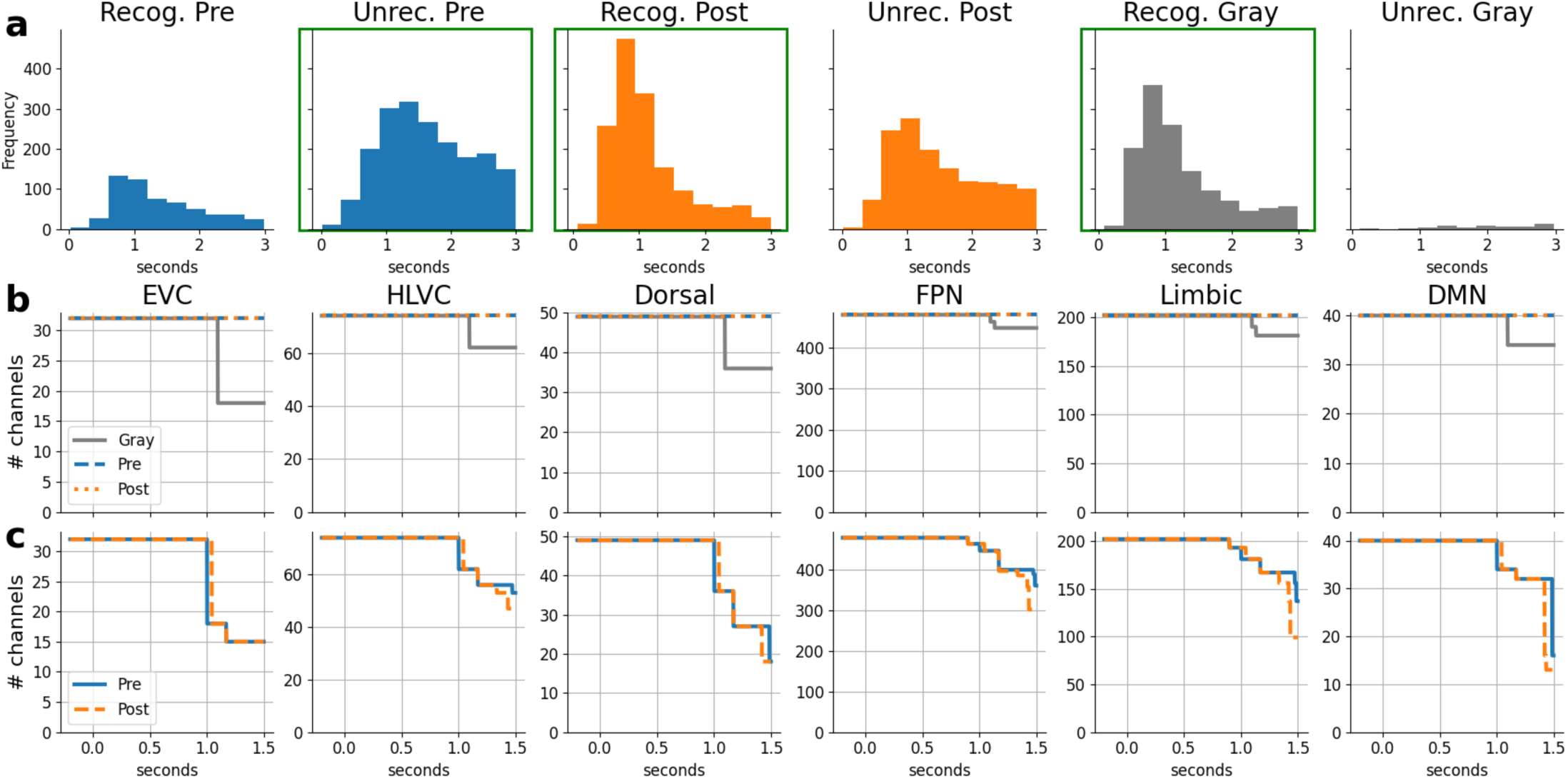
Reaction time distribution and trial number statistics for the iEEG experiment. **(a)** Histograms of trial-level reaction times, pooled across all subjects. Reaction time is the duration from image onset to the subject’s keyboard button press response. Trials are defined as ‘Recognized’ if it had a ‘Yes’ keyboard response and a correct verbal response (for details, see Methods, iEEG Experimental Paradigm). Trials with no keyboard response within the 3 sec period are not included. Unrecognized Pre, Recognized Post, and Recognized Gray trials (outlined in green) are used in the mean ROI activation and Image Preference analyses. **(b)** Electrode survival plots across trial epoch for mean ROI activation analysis. An electrode drops out of the analysis once all trials of the given type have ended, corresponding to the patient providing all keyboard responses prior to that timepoint. **(c)** As **b** for Image Preference analysis. An electrode drops out once a valid slope cannot be calculated for the trial type; see Methods, Image Preference Analysis. For trial type Pre (or Post), a valid slope calculation requires at least two images that each have at least one Grayscale trial and one Pre (or Post) trial remaining (i.e., the keyboard response happens after that timepoint).

**Supplementary Fig. 8.**
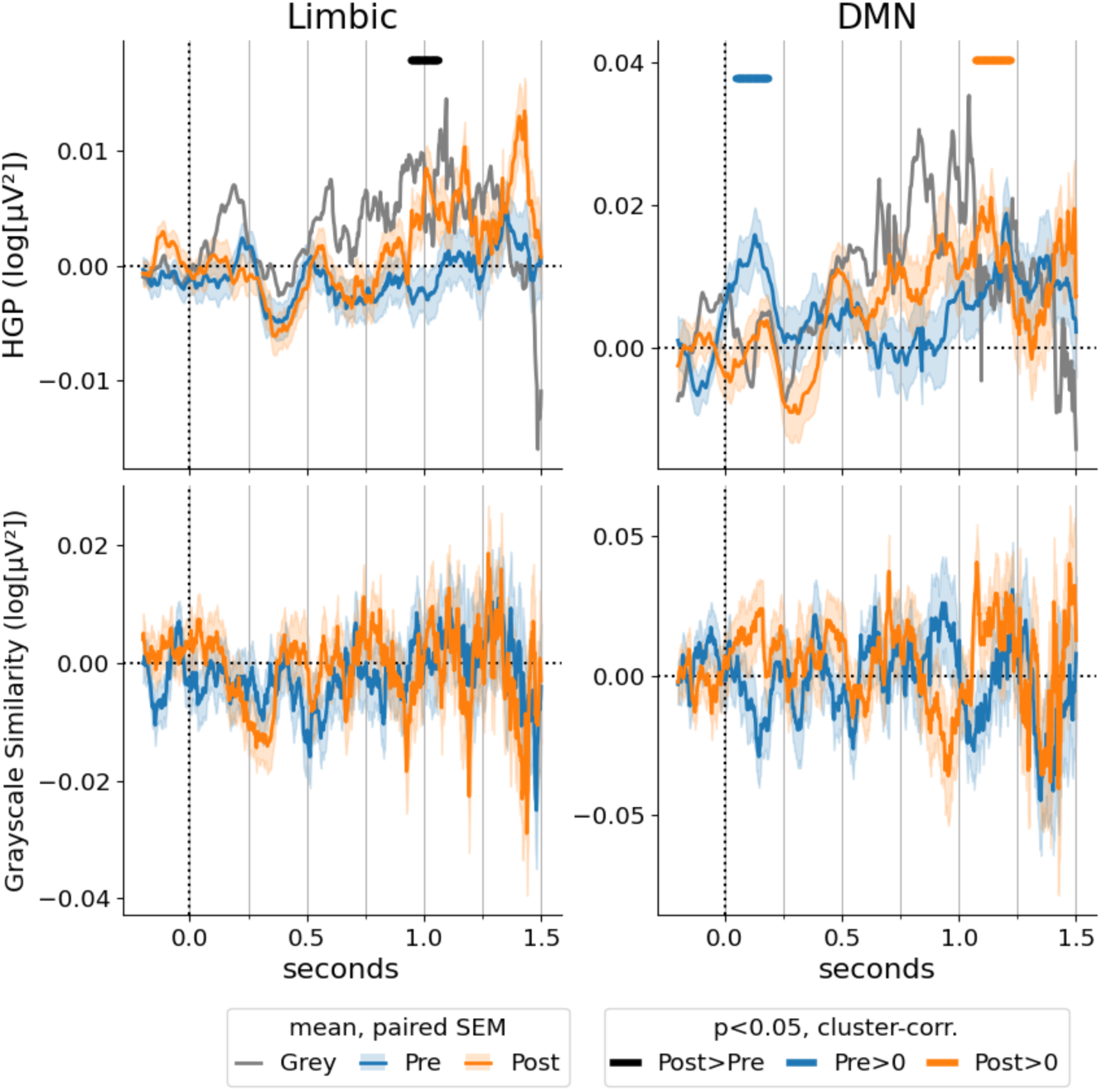
iEEG results for additional ROIs (limbic network and default-mode network). Figure layout is the same as main text Figure 4c.

**Supplementary Fig. 9.**
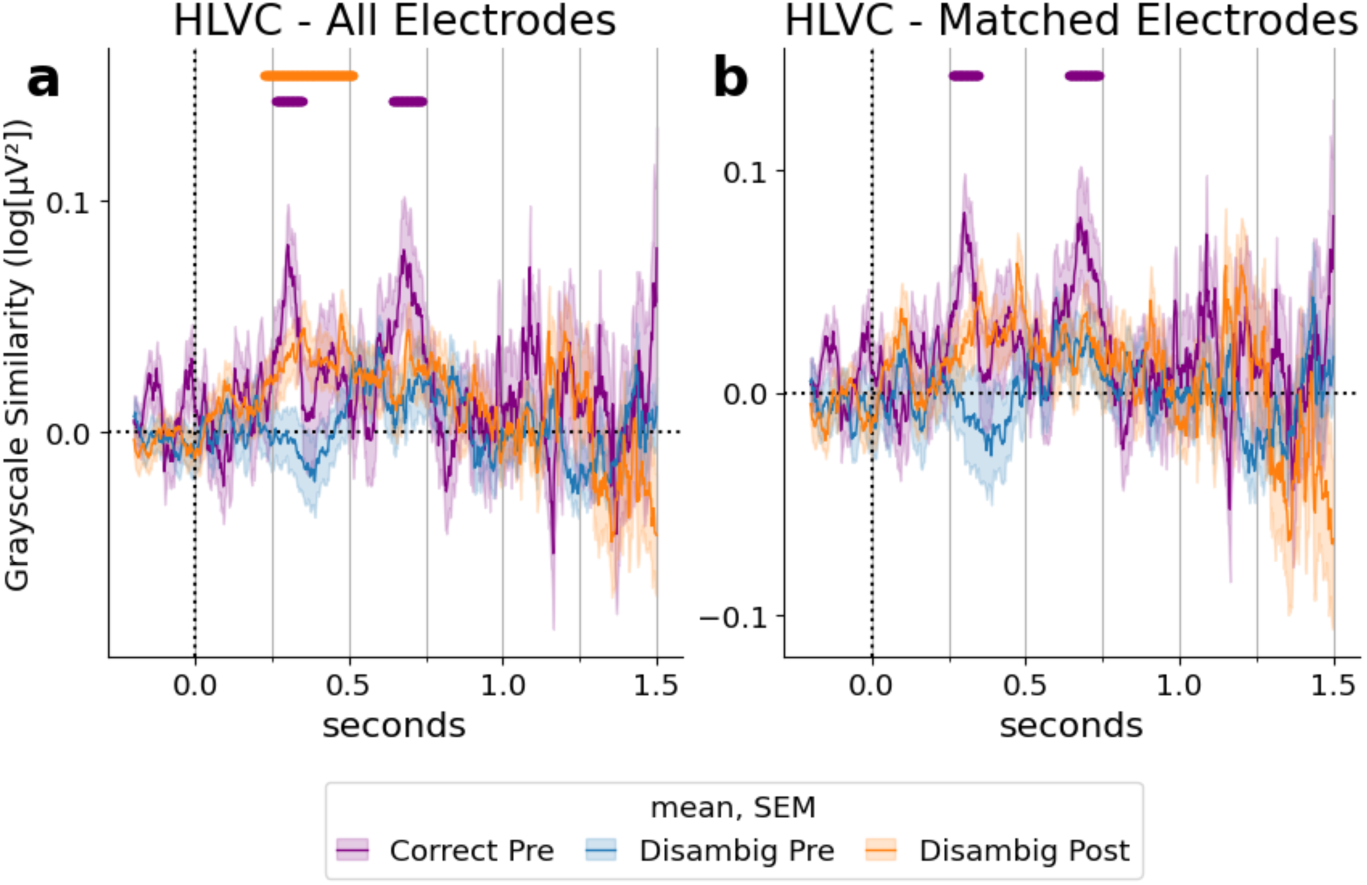
Greyscale Similarity for additional trial types in HLVC. Blue and orange traces are computed using pre- and post-Mooney images exhibiting the classic disambiguation effect; i.e., images used in the analysis of Fig. 4. Purple traces are from spontaneously recognized pre-Mooney images. Because pre-Mooney images had a low recognition rate (see Supplementary Fig. 6c, left), only 46 electrodes in HLVC could be analyzed for this analysis (two patients had insufficient correct pre trials), instead of the original 74 electrodes used in the analysis of Fig. 4. (**a**) Grayscale tuning similarity time course for pre-recognized images (purple) obtained using these 46 electrodes is plotted together with the original image-preference analysis result for HLVC obtained using 74 electrodes (blue and orange, identical to Fig. 4c, bottom, HLVC plot). **(b)** Same as (a), but the disambiguated pre and post results (blue and orange) were recalculated using the same 46 electrodes as used with the correct pre condition. Significance bars: p<0.05, cluster-based permutation test (based on Wilcoxon sign-rank test). Traces are mean similarity across electrodes, with shaded areas corresponding to SEM of the corresponding condition.

**Supplementary Fig. 10.**
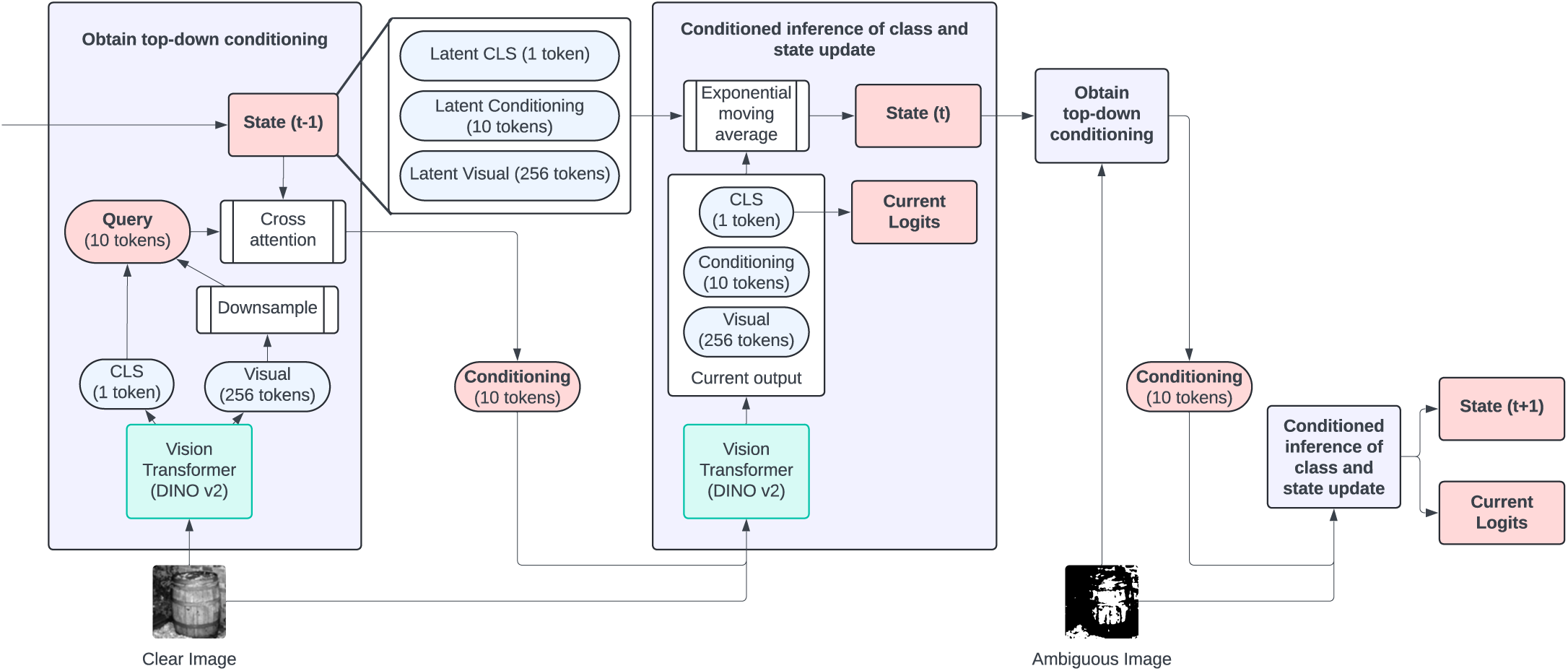
Detailed figure of our model architecture. For detailed description, refer to methods.

**Supplementary Fig. 11.**
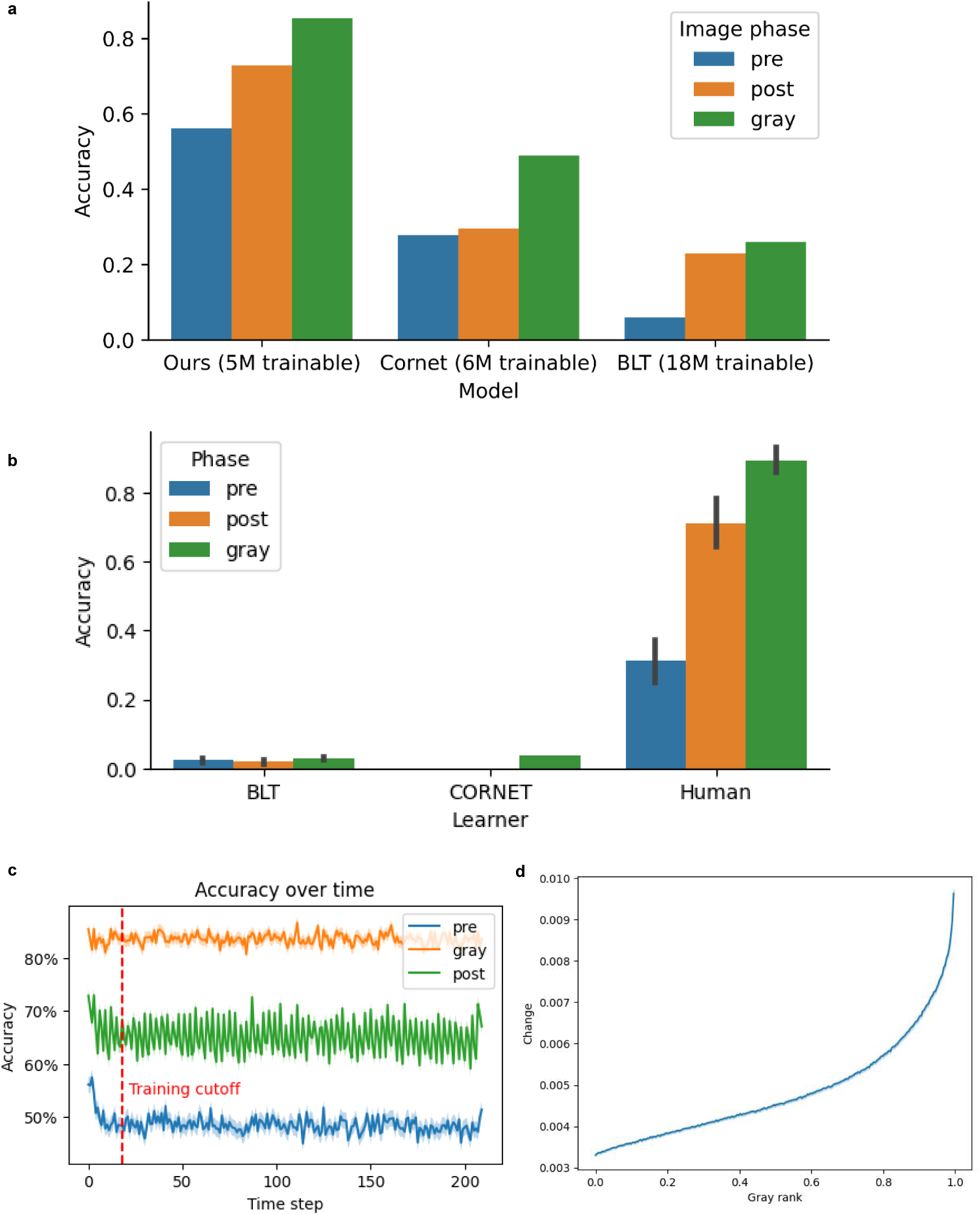
Detailed model evaluation. (**a**) Model performance comparison with existing neuroscience-inspired vision models. Performance is evaluated using 1,000 image sequences following the same format as those used for training the model but created using a held-out set of ImageNet1k images. (**b**) Although BLT and CORNet performs above chance on the held-out image sequences following the training image sequence format (in a), they lose their performance when evaluated on the same sequences of images used in the human experiment that are much longer. The performance of our novel DNN model on the same image sequences is shown in Fig. 5c. Error bars show 95% CI over subjects. (**c**) Our model’s long sequence performance. Model shows consistent learning performance when evaluated on Mooney learning sequences much longer than training (red dash line). Plot is generated by averaging over 10,000 randomly generated sequences using images from ImageNet1k. Shaded area shows 95% CI over trials. (**d**) Image patches with a higher attention rank, measuring its relative perceptual importance, during the disambiguation phase (i.e., grayscale image input) have greater changes in attention score from pre phase to post phase.

**Supplementary Fig. 12.**
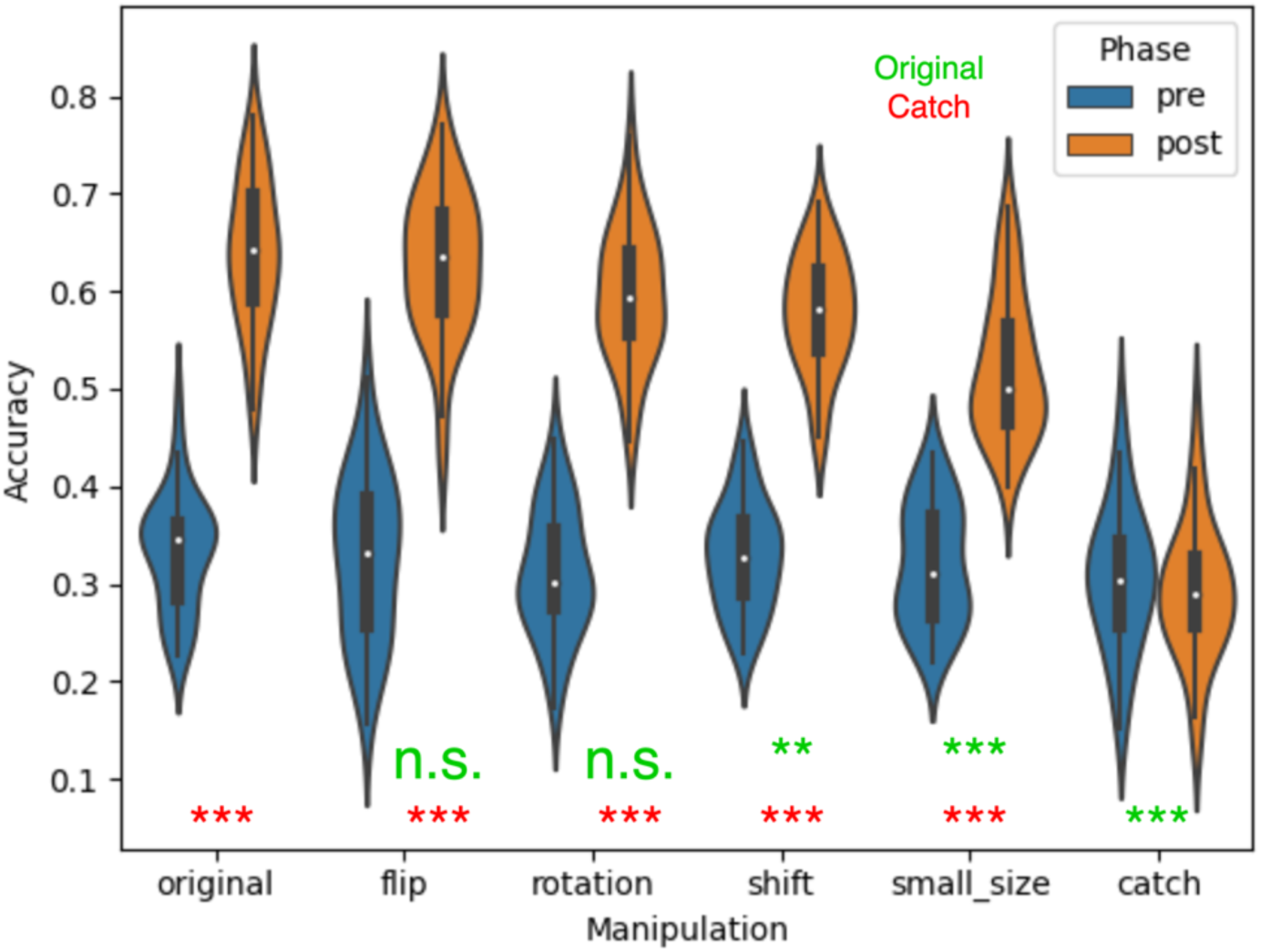
Model behavioral invariance analysis. Model behavioral invariances when evaluated on Mooney image trials with manipulations applied to grayscale images, analogous to the human experiment shown in Fig. 2a-e. Green and red asterisks refer to the interaction effect in a 2×2 ANOVA ([pre vs. post] x [manipulation vs. original/catch]) comparing each experimental condition with the original and catch condition, respectively. **: p < 0.01; ***: p < 0.001. Our model shows partial behavioral invariance to position shift and small size manipulation (interaction effect compared to original: p=5.10×10^-3^, p=6.70×10^-6^) and full invariance to 90-degree rotation and horizontal flip (interaction effect compared to original: p=0.792, p=0.239). All manipulations remain highly significant when compared to catch trials (all ps < 0.001). Evaluated using 20 random trials identical to the online pilot study, but with random manipulation applied to the grayscale image.

**Supplementary Fig. 13.**
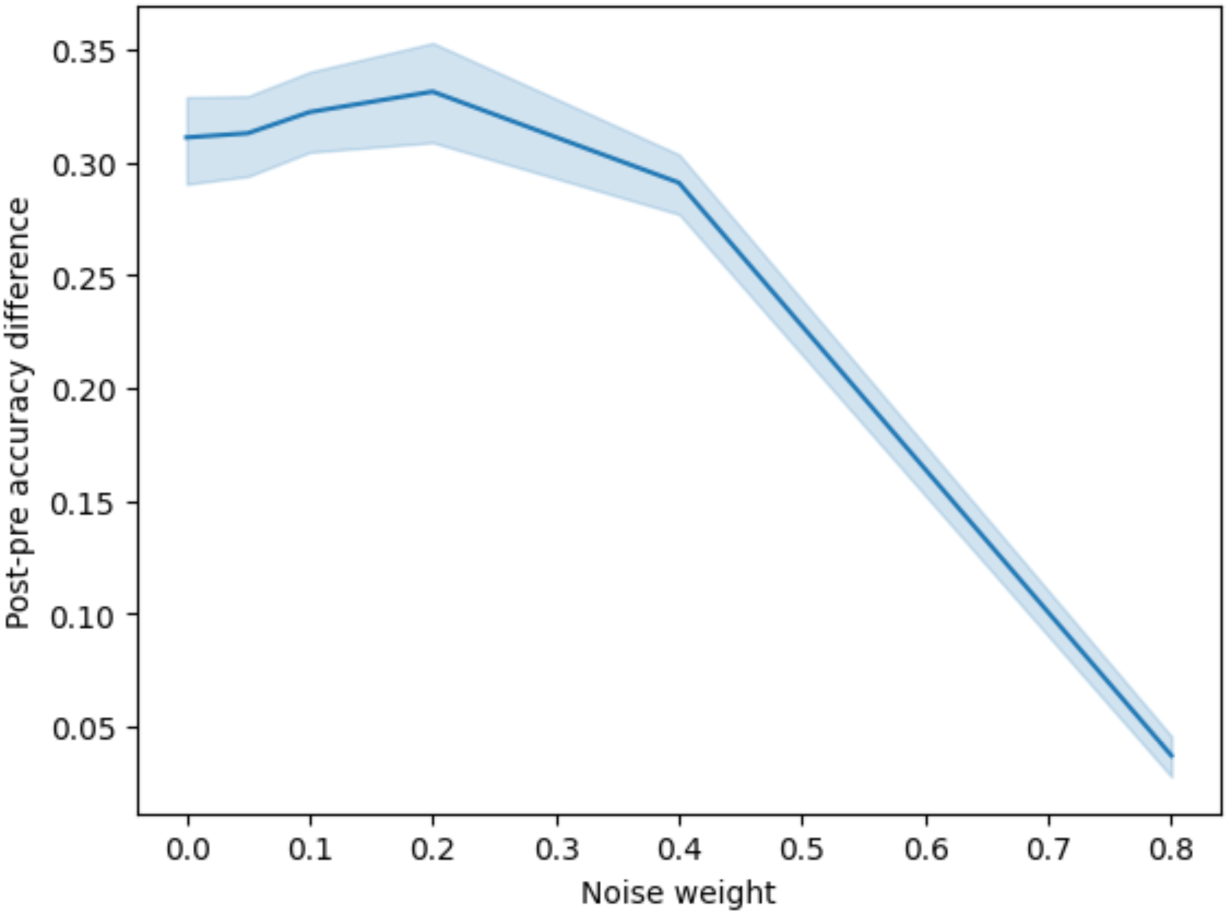
Model accuracy degradation with corrupted conditioning signal. We linearly mix the conditioning signal with norm-matched Gaussian noise using a weighted average operation. Result was evaluated with weight = [0.05, 0.1, 0.2, 0.4, 0.8]. The difference between post-phase accuracy and pre-phase accuracy decreases as the conditioning signal is corrupted by random norm-matched noise. Shaded area shows 95% CI over trials.

## Supplementary Tables

**Supplementary Table 1:**
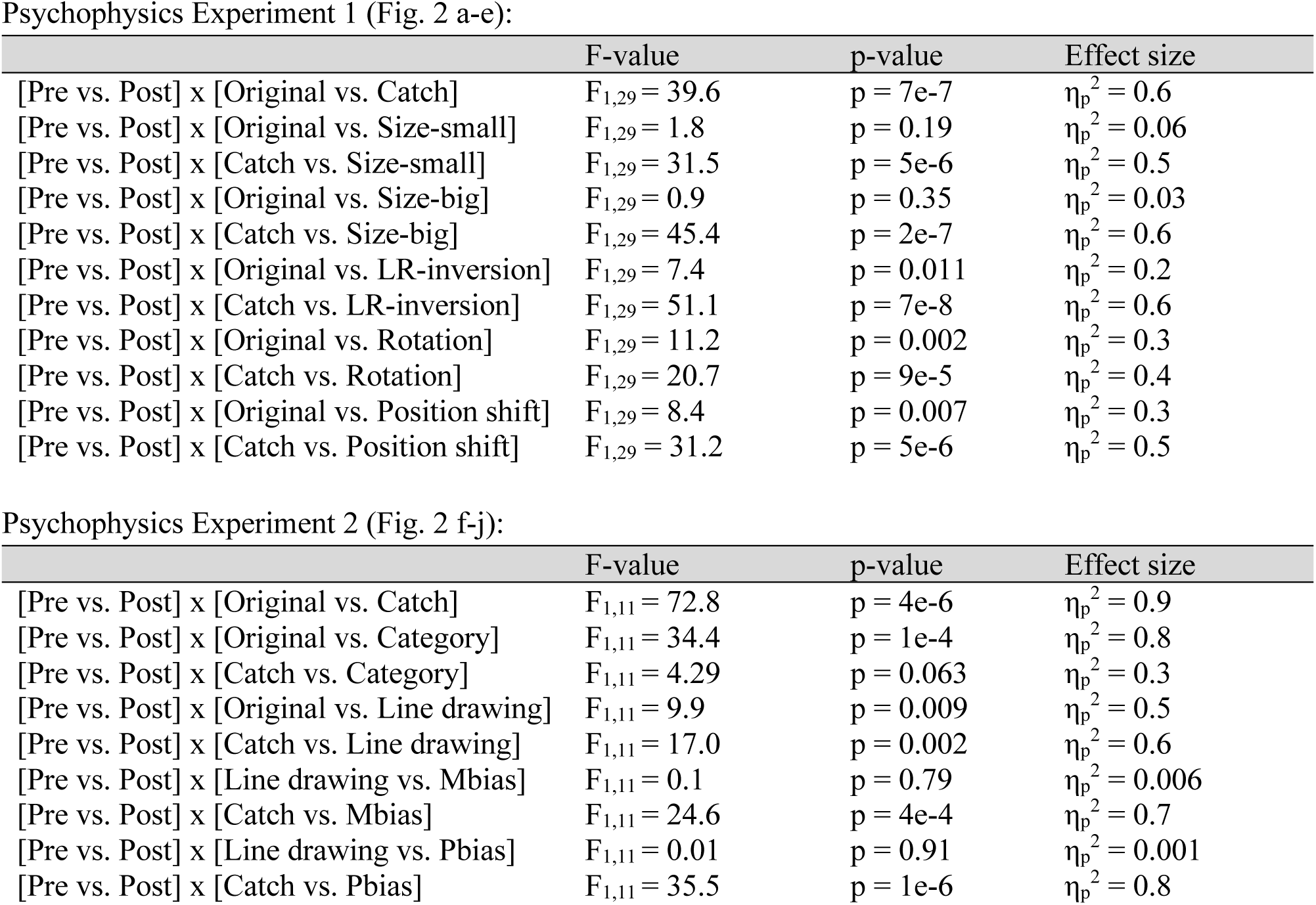
Full statistics for the psychophysics experiment. Top and bottom describe statistics related to Experiment 1 and Experiment 2, respectively, including effect sizes that are not reported in the main text. Only interaction effects in the two-way repeated measures ANOVA are shown, for brevity. Note that data from Experiment 1, original and catch condition, are plotted in both Fig. 2a and Fig. 1d.

**Supplementary Table 2:**
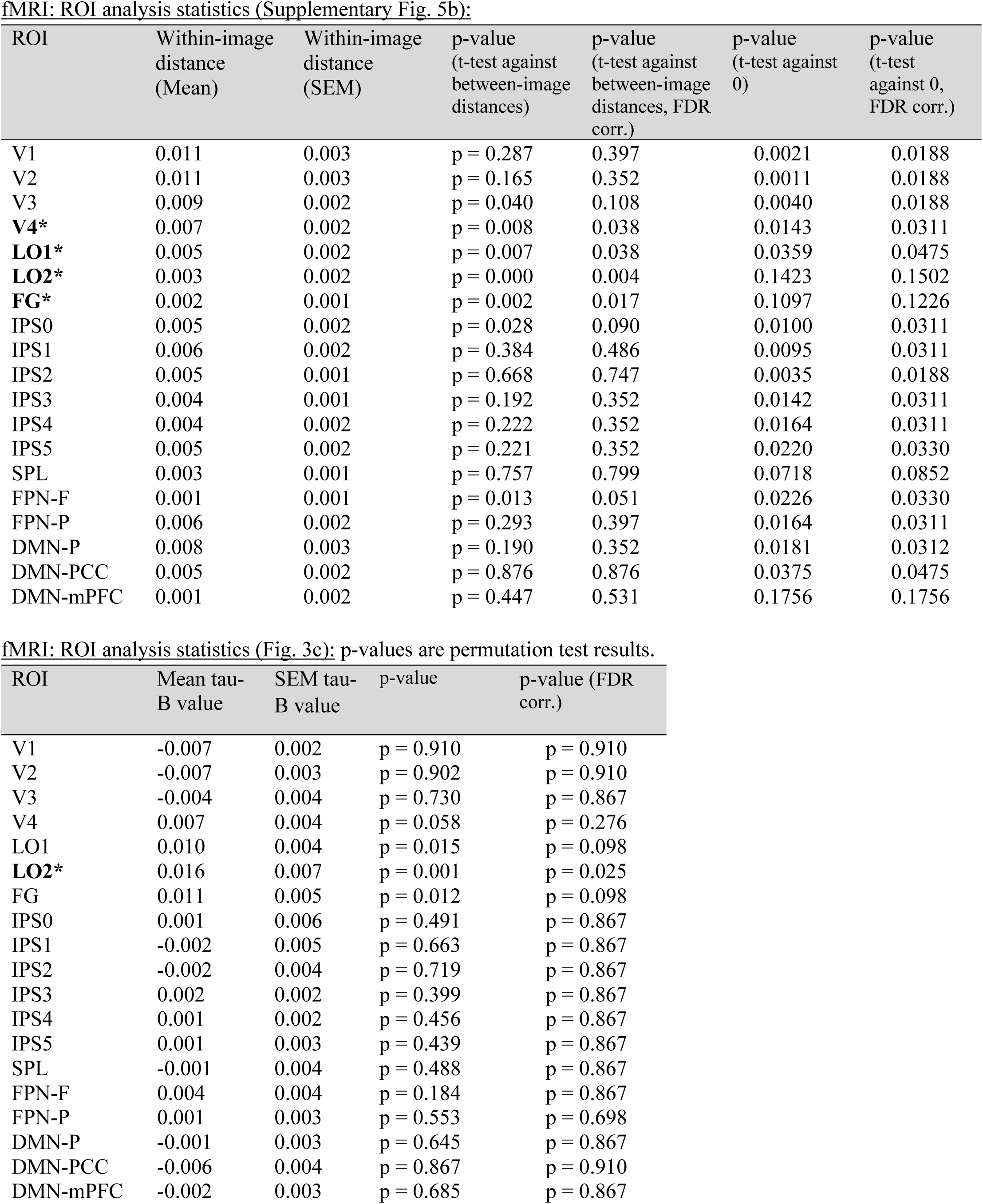
Full statistical results accompanying Supplementary Fig. 5b (top) and Fig. 3c (bottom). Top: statistical test evaluated the mean within-image distances (column 2) against a distribution of subsampled between-image distances (p-values in columns 4 and 5) and one-sample t-test against 0 (p-values in columns 6 and 7). Bottom: statistical test evaluated Kendall’s tau values against a null distribution obtained from shuffled image labels. FDR correction was applied across all 19 ROIs for each analysis.

| ROI | Within-image distance (Mean) | Within-image distance (SEM) | p-value (t-test against between-image distances) | p-value (t-test against between-image distances, FDR corr.) | p-value (t-test against 0) | p-value (t-test against 0, FDR corr.) |
| --- | --- | --- | --- | --- | --- | --- |
| V1 | 0.011 | 0.003 | p = 0.287 | 0.397 | 0.0021 | 0.0188 |
| V2 | 0.011 | 0.003 | p = 0.165 | 0.352 | 0.0011 | 0.0188 |
| V3 | 0.009 | 0.002 | p = 0.040 | 0.108 | 0.0040 | 0.0188 |
| **V4\*** | 0.007 | 0.002 | p = 0.008 | 0.038 | 0.0143 | 0.0311 |
| **LO1\*** | 0.005 | 0.002 | p = 0.007 | 0.038 | 0.0359 | 0.0475 |
| **LO2\*** | 0.003 | 0.002 | p = 0.000 | 0.004 | 0.1423 | 0.1502 |
| **FG\*** | 0.002 | 0.001 | p = 0.002 | 0.017 | 0.1097 | 0.1226 |
| IPS0 | 0.005 | 0.002 | p = 0.028 | 0.090 | 0.0100 | 0.0311 |
| IPS1 | 0.006 | 0.002 | p = 0.384 | 0.486 | 0.0095 | 0.0311 |
| IPS2 | 0.005 | 0.001 | p = 0.668 | 0.747 | 0.0035 | 0.0188 |
| IPS3 | 0.004 | 0.001 | p = 0.192 | 0.352 | 0.0142 | 0.0311 |
| IPS4 | 0.004 | 0.002 | p = 0.222 | 0.352 | 0.0164 | 0.0311 |
| IPS5 | 0.005 | 0.002 | p = 0.221 | 0.352 | 0.0220 | 0.0330 |
| SPL | 0.003 | 0.001 | p = 0.757 | 0.799 | 0.0718 | 0.0852 |
| FPN-F | 0.001 | 0.001 | p = 0.013 | 0.051 | 0.0226 | 0.0330 |
| FPN-P | 0.006 | 0.002 | p = 0.293 | 0.397 | 0.0164 | 0.0311 |
| DMN-P | 0.008 | 0.003 | p = 0.190 | 0.352 | 0.0181 | 0.0312 |
| DMN-PCC | 0.005 | 0.002 | p = 0.876 | 0.876 | 0.0375 | 0.0475 |
| DMN-mPFC | 0.001 | 0.002 | p = 0.447 | 0.531 | 0.1756 | 0.1756 |

**Supplementary Table 2: Full statistical results accompanying Supplementary Fig.**
| ROI | Mean tau-B value | SEM tau-B value | p-value | p-value (FDR corr.) |
| --- | --- | --- | --- | --- |
| V1 | -0.007 | 0.002 | p = 0.910 | p = 0.910 |
| V2 | -0.007 | 0.003 | p = 0.902 | p = 0.910 |
| V3 | -0.004 | 0.004 | p = 0.730 | p = 0.867 |
| V4 | 0.007 | 0.004 | p = 0.058 | p = 0.276 |
| LO1 | 0.010 | 0.004 | p = 0.015 | p = 0.098 |
| **LO2\*** | 0.016 | 0.007 | p = 0.001 | p = 0.025 |
| FG | 0.011 | 0.005 | p = 0.012 | p = 0.098 |
| IPS0 | 0.001 | 0.006 | p = 0.491 | p = 0.867 |
| IPS1 | -0.002 | 0.005 | p = 0.663 | p = 0.867 |
| IPS2 | -0.002 | 0.004 | p = 0.719 | p = 0.867 |
| IPS3 | 0.002 | 0.002 | p = 0.399 | p = 0.867 |
| IPS4 | 0.001 | 0.002 | p = 0.456 | p = 0.867 |
| IPS5 | 0.001 | 0.003 | p = 0.439 | p = 0.867 |
| SPL | -0.001 | 0.004 | p = 0.488 | p = 0.867 |
| FPN-F | 0.004 | 0.004 | p = 0.184 | p = 0.867 |
| FPN-P | 0.001 | 0.003 | p = 0.553 | p = 0.698 |
| DMN-P | -0.001 | 0.003 | p = 0.645 | p = 0.867 |
| DMN-PCC | -0.006 | 0.004 | p = 0.867 | p = 0.910 |
| DMN-mPFC | -0.002 | 0.003 | p = 0.685 | p = 0.867 |

**Supplementary Table 3:** Electrode count details. “All recorded” includes all electrodes that were not excluded prior to any analysis. Exclusion causes of this type included out of brain location, in lesion location, pervasive artifacts identified during preprocessing, and poor contact. “All analyzed” is a subset of all recorded electrodes that were assigned to an ROI and included in the analyses. Subsequent rows indicate how many electrodes were members of each network ROI and numbers of electrodes residing in multiple network ROIs. Because different network ROIs had overlaps, an electrode could be simultaneously assigned to multiple network ROIs. See Methods and Supplementary Fig. 4 for detailed ROI definitions and locations.

|  |  | Electrode Counts |
| --- | --- | --- |
|  | All recorded | 1886 |
|  | All analyzed (within ROIs) | 732 |
| Electrode counts for each network ROI | FPN | 479 |
|  | Limbic | 202 |
|  | HLVC | 74 |
|  | Dorsal | 49 |
|  | DMN | 40 |
|  | EVC | 32 |
| Number of electrodes residing in more than one network ROIs | FPN and Limbic | 83 |
|  | Dorsal Stream and FPN | 22 |
|  | DMN and FPN | 19 |
|  | EVC and HLVC | 14 |
|  | DMN and Dorsal Stream | 11 |
|  | Dorsal Stream and HLVC | 3 |
|  | DMN and EVC | 1 |
|  | DMN and HLVC | 1 |
|  | DMN and Dorsal Stream and FPN | 10 |

**Supplementary Table 4:** Demographic, clinical and data collection for each iEEG patient. Patients’ age at surgery ranged from 22 to 71 years old (mean: 35.7; std = 13.0). 16 (/three) out of 19 patients were females (/males). Seizure type: FAS = Focal aware seizure, FIAS = Focal with impaired awareness seizure, F2BTC = Focal to bilateral tonic-clonic seizure. Seizure focus: **bolded** entries indicate patients for whom resection, laser ablation, or (in one case, frontal) disconnection was performed, in which case the resection/ablation/disconnection sites were reported. When resection/ablation was not carried out due to multifocal seizures or ictal onset zones overlapping with eloquent cortex, the ictal onset zones are reported (most of these patients received RNS treatment). Electrode coverage information for each patient can be found in Supplementary Fig. 6. # electrodes recorded: number of electrodes that were not excluded prior to any analysis (see Supplementary Table 3 caption for more details). # electrodes analyzed: subset of all recorded electrodes that were assigned to an ROI and included in the analyses reported in Fig. 4 and Supplementary Fig. 8.

| # | Recording Center | Handedness | Seizure Type | Seizure Focus | # electrodes recorded | # electrodes analyzed | # Runs Recorded / Analyzed |
| --- | --- | --- | --- | --- | --- | --- | --- |
| 1 | NYU | L | FIAS, F2BTC | **right orbitofrontal cortex** | 151 | 69 | 14/14 |
| 2 | NYU | R | FAS, F2BTC | left parahippocampal gyrus, left insula | 164 | 58 | 14/14 |
| 3 | NYU | R | FIAS | **right subfrontal neocortex, right anterior temporal cortex, right hippocampus** | 77 | 15 | 13/13 |
| 4 | NYU | L | F2BTC, FIAS, FAS | **right anterior temporal neocortex, right hippocampus** | 68 | 35 | 14/14 |
| 5 | NYU | R | F2BTC, FIAS, FAS | **right anterior temporal neocortex, right hippocampus** | 113 | 45 | 14/14 |
| 6 | Mt Sinai | R | FAS, FIAS | **left hippocampus** | 70 | 22 | 14/14 |
| 7 | NYU | R | FAS | left anterior insula | 137 | 65 | 14/14 |
| 8 | NYU | R | FAS, FIAS | left amygdala, hippocampus, parahippocampal gyrus, temporal pole | 66 | 18 | 14/14 |
| 9 | Mt Sinai | R | F2BTC, FIAS, FAS | **right frontal (TBI)** | 70 | 31 | 14/13 |
| 10 | Mt Sinai | R | FAS | left motor strip | 53 | 11 | 14/14 |
| 11 | Mt Sinai | R | F2BTC, FIAS, FAS | left hippocampus, left post-lateral temporal | 65 | 31 | 12/12 |
| 12 | Mt Sinai | R | F2BTC, FIAS | left lateral temporal, left frontal operculum | 70 | 23 | 14/14 |
| 13 | NYU | R | FIAS, F2BTC | bilateral parietal occipital | 136 | 59 | 14/14 |
| 14 | Mt Sinai | R | FIAS | **right hippocampus** | 118 | 49 | 14/14 |
| 15 | NYU | R | FIAS, F2BTC | left mesial temporal/temporal pole | 75 | 15 | 14/14 |
| 16 | NYU | R | F2BTC | right anteromesial temporal (peri-amygdala region) | 84 | 36 | 14/14 |
| 17 | Mt Sinai | R | F2BTC, FIAS, FAS | bilateral hippocampi, right orbitofrontal | 113 | 48 | 14/14 |
| 18 | NYU | R | F2BTC, FIAS, FAS | Right temporo-occipital junction, superior parietal cortex, posterior dorsal insula | 96 | 52 | 14/14 |
| 19 | NYU | R | FIAS | left mid-temporal and posterior temporal, insular | 160 | 50 | 14/14 |

## Supplementary Methods

### Online Pilot Experiment

#### Subjects

26 subjects were recruited from the Prolific.co participant pool, but 10 subjects were rejected due to not using a desktop monitor, and 4 for timing out of the study. Data from 12 subjects were used to select the top 90 Mooney images used in the in-person behavioral experiment. Gender and age information were not collected; however, participants confirmed right-handedness and normal or corrected-to-normal vision. All participants were provided with a written informed consent, and the experiment was approved by the Institutional Review Board of New York University School of Medicine (protocol #S21-00218).

#### Experimental Stimuli

The task was created using Gorilla.sc, an online behavior experiment platform. Screen size was individually calibrated for each participant on their desktop monitor, using a credit card, and participants were instructed to maintain a consistent viewing distance from the screen. 219 images were taken from Caltech and Pascal VOC grayscale image databases that contain real-world objects and animals. We followed a previous protocol to generate Mooney images^5^. Briefly, the images were converted to grayscale and fed through a low-pass filter (a 2D Gaussian smoothing kernel with a standard deviation of 1.5) and binarized to create the two-tone Mooney images.

#### Experimental procedure

First, participants briefly ran practice trials with and without time limits. The main task consisted of a single session divided into 3 task blocks, and median completion time was 75 minutes. 219 Pre-learning Mooney images, and corresponding grayscale and Post-learning Mooney images were shown in block-shuffled order as described in Fig. 1a. In each trial, fixation cross was presented for 1 second, followed by an image presentation (Mooney or grayscale) for up to 2 seconds until text-entry response. The text input box remained on the screen for up to 6 seconds. Participants typed their responses using their keyboard into a text input box. Instructions specified to provide exemplar label (e.g., “cat’), not broad category labels (e.g., “animal”).

#### Analysis

All data were extracted from Gorilla.sc and analyzed in MATLAB using custom-written scripts. An automated script compared typed responses and a pre-generated answer key, and were later manually inspected to include spelling errors. Category responses (e.g., “animal” instead of “dog”) were marked incorrect. Images were then scored based on an optimal-learning criteria, where the Mooney image is initially unrecognized, followed by recognition after grayscale presentation. Then, images were ranked based on the similarity of optimal learning criteria across subjects. The best 90 images, defined by the largest performance improvements from the pre to post stage, were chosen to be used in the in-person behavioral experiment.

### Behavioral Experiment 1

#### Subjects

33 participants were recruited from the greater New York City area. Ages 20-70 (median age 28, std = 13.7), 18 were female. Most of the participants (31 out of 33) were right-handed, and their vision was normal or corrected-to-normal. All participants were provided with a written informed consent, and the experiment was approved by the Institutional Review Board of New York University School of Medicine (protocol #S15-01323).

Complete data from 3 participants were excluded due poor performance in the main task. 1 block was removed from 2 subjects due to experimental script error and request to leave early, respectively. Exclusion criteria was established prior to the beginning of the study.

Data from a total of 30 participants were used in the final analysis.

#### Experimental Stimuli

The task was created using PsychoPy 2020.1.3 and presented on a 1920 × 1080 monitor, placed 63 cm away from the participant’s eyes. Participants placed their heads on a chin rest to minimize head movements and ensure consistent viewing angle. In the original trials, the Mooney and grayscale image had the same retinal location, size, and orientation (12 dva in size, presented at central fixation).

#### Experimental Procedure

Participants first completed two blocks of practice trials, first without a time limit for task familiarity, and later with time limits used in the main task. In each trial, purple fixation dot was presented for 1 second, followed by an image presentation (Mooney or grayscale) for 2 seconds. Subjects responded with a Yes/No recognition button press, followed by a verbal response (with an upper limit of 6 seconds), which was recorded in real-time. Participants completed the task inside a dimly lit, soundproof room designed for EEG studies; a microphone was fed through the cable mount so that verbal responses could be heard from outside the room. Eye-tracking data were recorded using EyeLink 1000, in the binocular mode with a sampling rate of 1000 Hz.

Main task consisted of 3 blocks of 90 trials each. In total, 90 unique Mooney images were assessed. The following grayscale image conditions were tested: original, catch, size-small (6 dva), size-large (24 dva), visual field shift (6 dva left/right shift), left-right inversion, 90° rotation (CW/CCW). Each unique Mooney image was assigned to a single condition for each subject (because each image can only be tested once/participant), and this assignment, as well as the presentation order of conditions were counterbalanced across participants. The counter-balance structure is shown in the table below. M1-M8 denotes the 7 image manipulation conditions listed above, plus an additional condition not relevant to the present study. Thirty participants were included in this experiment, and were evenly distributed across the 10 groups (i.e., each group contained 3 subjects). All images were randomly shuffled across all manipulation groups, to prevent trial blocks of the same manipulation. Results were pooled and averaged across all subjects to test learning outcomes for each condition. This counter-balancing design was necessary because each participant can only view each unique image in one condition, given the one-shot perceptual learning task.

**Experiment 1: Counterbalanced groups**
| Images:<br>1-9 | Images:<br>10-18 | Images:<br>19-27 | Images:<br>28-36 | Images:<br>37-45 | Images:<br>46-54 | Images:<br>55-63 | Images:<br>64-72 | Images:<br>73-81 | Images:<br>82-90 |  |
| --- | --- | --- | --- | --- | --- | --- | --- | --- | --- | --- |
| M1 | M2 | M3 | M4 | M5 | M6 | M7 | M8 | Original | Catch | Group 1 |
| Catch | M1 | M2 | M3 | M4 | M5 | M6 | M7 | M8 | Original | Group 2 |
| Original | Catch | M1 | M2 | M3 | M4 | M5 | M6 | M7 | M8 | Group 3 |
| M8 | Original | Catch | M1 | M2 | M3 | M4 | M5 | M6 | M7 | Group 4 |
| M7 | M8 | Original | Catch | M1 | M2 | M3 | M4 | M5 | M6 | Group 5 |
| M6 | M7 | M8 | Original | Catch | M1 | M2 | M3 | M4 | M5 | Group 6 |
| M5 | M6 | M7 | M8 | Original | Catch | M1 | M2 | M3 | M4 | Group 7 |
| M4 | M5 | M6 | M7 | M8 | Original | Catch | M1 | M2 | M3 | Group 8 |
| M3 | M4 | M5 | M6 | M7 | M8 | Original | Catch | M1 | M2 | Group 9 |
| M2 | M3 | M4 | M5 | M6 | M7 | M8 | Original | Catch | M1 | Group 10 |

Following previous studies^4, 6^, images were presented in mini-blocks that consisted of 3 grayscale images followed by 6 Mooney images. The 6 Mooney images included 3 “Post” images that correspond to the 3 grayscale images shown just before, and 3 “Pre” images that correspond to grayscale images that would be shown in the subsequent block, and their order were randomly shuffled. Before the first block, 3 “Pre” images were shown before the start of this block structure. For the last block, only 3 grayscale images and 3 “Post” images were shown.

### Behavioral Experiment 2

#### Subjects

12 participants were recruited from the greater New York City area. Ages 20-61 (median age 26.5, std = 14.5), 8 were female. Vision was normal or corrected-to-normal, and 10 out of 12 participants were right-handed). All participants were provided with written informed consent (protocol #S15-01323). Data from all 12 participants were used in the final analysis.

#### Experimental Stimuli

Task structure and experimental set-up is similar to Experiment 1, except that a different set of grayscale image manipulation conditions were used. In addition to original and catch trials, which were identical to those in Experiment 1, grayscale images were manipulated in the following four conditions: category images (images within the same semantic category but containing a different exemplar, see Fig. 2g), line drawings, M-bias, and P-bias images. Line drawings are introduced in this experiment as a proper control for M-bias and P-bias drawings, which were modified versions of line drawings (low contrast and red-green isoluminance, respectively). The line drawings were created by hand-traced object boundaries in ImageJ to create an image with uniform contrast and luminance (Fig. 2h).

#### Experimental Procedure

The experiment consisted of a single session with 3 components (perceptual isoluminance and contrast calibration, followed by the main task). To create P-bias images, first, the red-green perceptual isoluminance level was determined for each subject using a heterochromatic flicker photometry task^7, 8^. A 3 dva radius circle was presented at fixation and flickered between red and green at 14Hz. Green was set at [0 140 0] and the participant adjusted the red luminance using a slider, until the flickering appeared to slow down or stop. This fusion effect indicated isoluminance between the red and green colors, or matched brightness. Then, the P-bias images were regenerated for every subject using the identified red luminance value.

For M-bias images, images were presented on a uniform gray background with light gray line drawings of sample objects. The image was set to a 3.5% contrast, according to previously reported protocols^7–9^. To this end, participants performed a QUEST adaptive staircase procedure^10^ with 3 interleaved staircases to determine the image contrast that yielded a 50% object recognition rate. In the staircase procedure, a line drawing of an animal not used in the main task were presented for each of the 3 interleaved staircases. Participants were instructed to report recognition of the whole animal (not just a glimpse). After the completion of the staircase, the reversal value (luminance value for a 50% recognition threshold) was identified, and the 3.5% contrast was then calculated using the reversal value as baseline. To set the contrast to 3.5%, the equation becomes: 0.035 = (L_new_ – L_reversal_) / L_reversal_ to calculate L_new_. Then, the L_new_ value was set for all line drawing luminance values in the M-bias condition during the session.

In the main task, participants still completed two blocks of practice trials and followed by the main task, with a similar block structure as in Experiment 1. The 90 unique Mooney images and their associated grayscale images were each assigned to one of 6 conditions for each subject (original, catch, category, line-drawing, M-bias, P-bias), with the assignment counterbalanced across subjects. The presentation order of different conditions was also counterbalanced across subjects, following the structure shown in the table below.

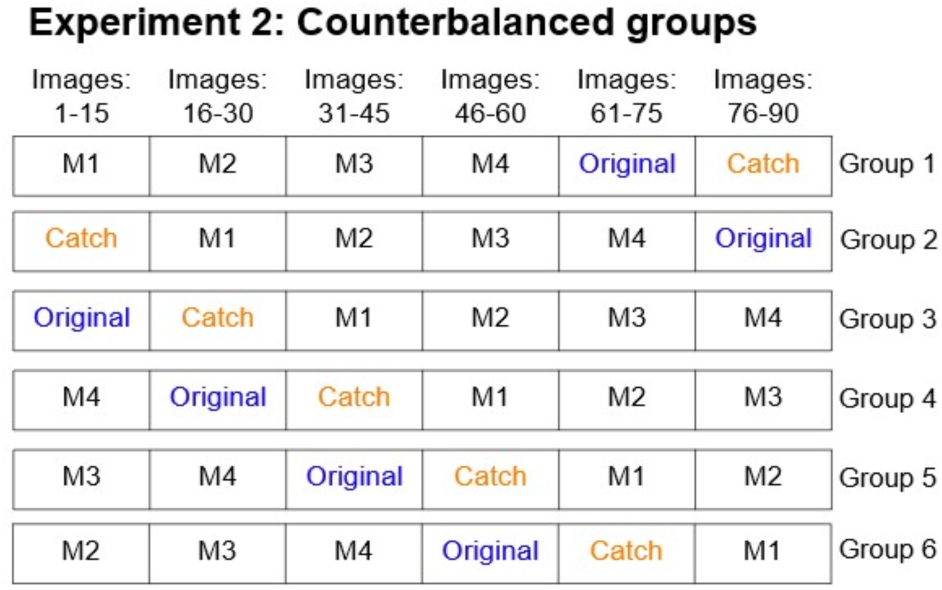

**Experiment 2: Counterbalanced groups**
| Images:<br>1-15 | Images:<br>16-30 | Images:<br>31-45 | Images:<br>46-60 | Images:<br>61-75 | Images:<br>76-90 |  |
| --- | --- | --- | --- | --- | --- | --- |
| M1 | M2 | M3 | M4 | Original | Catch | Group 1 |
| Catch | M1 | M2 | M3 | M4 | Original | Group 2 |
| Original | Catch | M1 | M2 | M3 | M4 | Group 3 |
| M4 | Original | Catch | M1 | M2 | M3 | Group 4 |
| M3 | M4 | Original | Catch | M1 | M2 | Group 5 |
| M2 | M3 | M4 | Original | Catch | M1 | Group 6 |

### Behavioral Experiment Data Analysis

#### Behavior data analysis

All data was extracted from PsychoPy and analyzed in MATLAB using custom-written scripts. Similar to the online behavior paradigm, responses were counted correct if they reported the correct exemplar, not a broad category (e.g., animal). For statistical testing, a two-way repeated measures ANOVA using both frequentist and Bayesian approaches, was conducted using Jeffreys’s Amazing Statistics Program (JASP)^11^.

The ANOVA was applied to the recognition responses to the Mooney images with the following factors: i) Mooney image presentation phase (Pre vs. Post); ii) grayscale image type (Original vs. Manipulated) or (Catch vs. Manipulated).

For Experiment 2, in the analysis of data from M-bias and P-bias image conditions, the original condition was replaced by line-drawing condition, such that the corresponding ANOVA had the following two factors: i) Mooney image presentation phase (Pre vs. Post); ii) grayscale image type (Line drawing vs. M-bias) or (Line drawing vs. P-bias).

#### Eye-tracking data analysis

The conversion from EDF files to a MATLAB compatible format was done with the Edf2Mat MATLAB Toolbox^12^, and further analysis was performed by custom MATLAB scripts. For VF shift trials, a control analysis was conducted in which trials with broken fixation were excluded (Supplementary Fig. 2). For this analysis, a threshold was applied to exclude any trials with a gaze shift outside of a 3-dva radius circle centered on fixation during stimulus presentation. Because of the uneven number of trials across conditions in this case, for statistics, a Generalized Linear Mixed Models (GLMM) with a logistic linking function was used, instead of an ANOVA, to model binary recognition outcomes across trials. This was implemented using the ‘fitglme’ function in MATLAB. In the model, the dependent variable is the recognition outcome; the fixed effect is the grayscale manipulation type, and random effects are subjects and trials. Interaction effects between Mooney image presentation phase (Pre vs. Post) and grayscale image type (Original vs. Manipulated) or (Catch vs. Manipulated) are reported.

### Population receptive field (pRF) Analysis

To quantify how much grayscale image manipulations may have altered neural coding at each stage of visual processing, we conducted a simulation using voxel-wise pRF data measured by fMRI in previous papers^1, 2^. Analyzed ROIs include: V1, V2, V3, hV4, LO1, LO2, inferior occipital gyrus (IOG), posterior fusiform gyrus (pFus), and mid-fusiform gyrus (mFus). LO1, LO2, IOG, pFus and mFus are regions within HLVC, arranged along the posterior-anterior axis, wherein size and orientation invariances were reported to emerge. For each voxel from each ROI, the data included the estimated center and size of its population receptive field. Only voxels in which the pRF model explained >50% variance (i.e., r^2^ > 0.5) were included in the analysis, which yielded the following number of analyzed pRFs for each ROI:

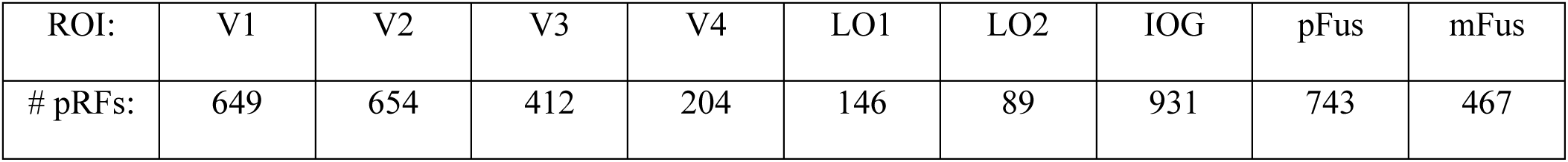

| ROI: | V1 | V2 | V3 | V4 | LO1 | LO2 | IOG | pFus | mFus |
| --- | --- | --- | --- | --- | --- | --- | --- | --- | --- |
| # pRFs: | 649 | 654 | 412 | 204 | 146 | 89 | 931 | 743 | 467 |

The analysis schematic is shown in Supplementary Fig. 1b. For every grayscale image, the most diagnostic image feature was traced by an investigator in Image J. For example, the defining image feature for the Dalmatian dog shown in the figure was labeled to be the face. Then, the image was binarized to create a black silhouette of the diagnostic image feature. For every ROI, all measured pRFs were overlayed with the diagnostic image feature for both the original and manipulated images. For each unique pRF, if it overlapped with at least 1 pixel in the image feature in the original image, the number of image feature pixels that overlapped with its pRF was counted, for both the original and manipulated images. In the example shown, the diagnostic image feature in the original image had 600 pixels that overlapped with the pRF of V1 voxel under investigation (outlined in red), and 0 pixels of the diagnostic image feature in the rotated image overlapped with the same voxel’s pRF.

Taking the ratio between these two values (# of pixels in the manipulated image that overlaps with the pRF / # of pixels in the original image that overlaps with the pRF), the pRF was labeled as “overlapping” if the ratio was at least 25%. This procedure was repeated for every voxel’s pRF, in every ROI, and separately for every manipulation. Finally, the number of pRFs that retained diagnostic image feature representation across the original-manipulation image pairs was counted, and the ratio between this number and the total number of measured pRFs in that ROI was computed. The result, showing the percentage of all measured pRFs that had significant overlap (>25%) between the original and manipulated images’ diagnostic feature, is plotted in Supplementary Fig. 1c. We acknowledge that the definition of diagnostic image feature was somewhat subjective, but this approach nevertheless allowed a quantitative estimate of the changes in neural coding at each level of ventral stream processing given a particular grayscale image manipulation.

### fMRI Experiment

#### Subjects

12 participants were recruited from the greater New York City area. Ages 21-42 (median age 23, std = 6.2), 9 were female, and all were right-handed with correct or corrected-to-normal vision. All participants were provided with a written informed consent, and the experiment was approved by the Institutional Review Board of New York University School of Medicine (protocol #S15-01323). Data from 2 participants were entirely excluded: one immediately opted out due to nausea in the scanner, and the other was excluded due to suboptimal scan quality. Lastly, 4 out of 16 blocks from 1 participant were excluded due to scanner error.

#### Experimental Stimuli

The task was created using PsychoPy 2021.2.3 and stimuli were presented using an MRI-compatible LCD monitor (BOLDScreen, Cambridge Research Systems) with a 120Hz refresh rate. The monitor was located 198 cm behind the center of the scanner bore, and participants viewed the screen using an eye mirror that was placed 5cm away from the participant’s eyes, attached to the head coil. To test for emergence of invariant object recognition, a subset of 10 grayscale images from the psychophysics study were used. The images were balanced between 5 animate and 5 inanimate objects. The images were shown in the following conditions that matched the psychophysics paradigm: Original image (11 dva), LR inversions, Rotation (CW), Rotation (CCW), Size-small (5.5 dva), VF shift (5.5 dva right), and VF shift (5.5 dva left), and line drawings. Line drawings were not further analyzed to constrain analysis to size, viewpoint, and position invariance from Experiment 1. Size-big (24 dva) manipulations were excluded due to monitor size and placement limitations inside the scanner room. Also, the size, and VF shift parameters had to be presented at a slightly smaller scale as compared to the behavioral experiments (from 12 to 11 dva, and from 6 to 5.5 dva) to accommodate scanner screen size.

#### Task design

First, participants were shown all possible 80 images (10 exemplars × 8 conditions) before entering the scanner (on a gray background, in similar format as the main task), for familiarity and to prevent any discrepancies in neural responses during the first and the subsequent runs. Each session consisted of anatomical scans and 16 runs of fMRI BOLD runs that were 5 minutes each, for a total of ∼90 minutes in the scanner. During the task inside the scanner, participants were asked to passively view the screen while maintaining visual fixation, and to respond with a button press when the fixation cross changed from white to red for a 200 ms duration. In each trial, the image was presented for 500ms, followed by a 1.5–3.5s jittered ITI. Each run included 80 trials, in which each unique image was presented once in shuffled order, with the constraint that two different manipulation conditions of the same image cannot be presented in adjacent trials (to avoid any repetition suppression/priming effects). The fixation cross color change happened 16 times per run, at a random time after the trial onset (during image presentation or the ITI). At the end of each run, subjects were given visual feedback about the proportion of successful button-presses in response to fixation cross color changes, to maintain task engagement.

#### MRI data acquisition

Experiments were run in a Siemens 7T MRI scanner using a 32-channel NOVA head coil at the NYU Center for Biomedical Imagining. T1 weighted MPRAGE images were acquired with 1.0 mm isotropic voxels, FOV 256 mm, 192 sagittal slices, TR 3000 ms, TE 4.49 ms, flip angle 6°, fat suppression on, bandwidth 130 Hz/Px. Proton density (PD) images were acquired for intensity normalization, with the following parameters: FOV 256 mm, 192 sagittal slices, 1.0 mm isotropic voxels, TR 1760 ms, TE 2.61 ms, flip angle 6°, bandwidth 280 Hz/Px. BOLD fMRI images were acquired using a GRE-EPI sequence with the following parameters: FOV 192 mm, 66 oblique slices covering all of cortex, voxel size 1.6 × 1.6mm, slice thickness 1.6mm with distance factor 10%, TR 1500 ms, TE 25 ms, multiband factor 2, GRAPPA acceleration 2, phase encoding direction posterior to anterior, flip angle 50°, bandwidth 1894 Hz/Px.

### fMRI Analysis

#### Preprocessing

Data preprocessing follows our published procedures^4, 13^. All fMRI analyses were preprocessed using FSL’s FEAT tool. Motion artifacts were corrected using MCFLIRT, which aligned each volume to the volume acquired in the middle of the run, and estimated 3 dimensions of head rotation and translation across time, with 6 DOF. Slice-timing correction accounted for the long whole-brain acquisition time of 1500ms, which interpolated the signals from each slice to the middle of each TR. Then, the brain was extracted using BET and spatial smoothing (3 mm FWHM) was applied. Lastly, ICA cleaning was used to remove artifacts related the motion, arteries, or CSF pulsation. The data was initially passed through AROMA ICA, an automatic artifact classification method, and 60-70 components that explain ∼80% of variance in the BOLD signal were manually inspected to select components that corresponded to artifacts. Functional images were registered to the individual subject’s MPRAGE (T1).

#### General linear model (GLM)

A general linear model (GLM) was used to extract stimulus-evoked activation, using the FEAT tool in FSL. For each task run, the following regressors were created: one regressor for each of the 80 unique images as well as the button press events, for a total of 81 regressors per run. For the button press regressor, a boxcar function was applied for the duration between the onset of fixation cross color change and the button press; in the event of missed trials, the boxcar lasted 200ms — the duration of color change. Then, beta estimates for each regressor were obtained. t-values were computed by dividing the beta estimate by its standard-error estimate (output from FSL), and were used for the rest of the analysis to suppress the contribution of noisy voxels in the beta estimate^14^. All analyses were conducted within each subject, with t-values aligned to the subject (T1) space.

#### Neural invariance analysis

Using the 70 × 70 neural RDM constructed by calculating cross-validated (c.v.) Euclidean distance between every pair of unique images (see main text Methods section, fMRI–Neural Distance Analysis), we plotted the mean within-image between-condition neural distances (green bars in Supplementary Fig. 5b). Two statistical analyses were performed using these values: 1) a statistical test for significant neural invariance by comparing within-image distances to across-image distances, and 2) an exploratory analysis involving t-tests against zero to test how far from “full invariance” each ROI is.

First, for each ROI, we tested whether there is significant neural invariance to image manipulations. Significant neural invariance would suggest that when the same image exemplar is shown under different manipulation conditions, the neural distance is significantly smaller than expected by chance. To calculate this chance level, we used between-image distances under the same image manipulation conditions. Specifically, for each ROI and each subject, the within-image across-condition neural distance was averaged across 210 values (21 condition pairs × 10 image exemplars). To create the null distribution (the expected upper-limit distance values for each ROI): 210 across-image distances were randomly drawn from the between-image part of the neural RDM (yellow region in the matrix in Supplementary Fig. 5a) for each subject, with equal distribution across condition pairs (i.e., matching the condition-pair distribution in the green bars). These values were averaged together and across all subjects to yield one sample of the null distribution, and this procedure was repeated 10,000 times to yield a null distribution for each ROI (yellow ribbon, Supplementary Fig. 5b). The within-image distances were then compared against this null distribution to obtain a p-value (one-tailed). Finally, FDR correction was applied across all ROIs to correct for multiple comparisons using the Benjamini-Hochberg method. Significant results indicate image-exemplar level representations that are invariant to image manipulations.

Second, to test how far the neural representation in an ROI is from full invariance, defined by no change in neural representation when the same image exemplar is manipulated, we tested the within-image between-condition neural distances against the chance level of 0 using a one-tailed t-test across subjects (FDR corrected). Importantly, c.v. Euclidean distance is an unbiased distance estimator, which is not conflated by noise (for details, see main text Methods section). Therefore, all measured distances should theoretically be independent of noise, and distances between patterns in response to identical images should be zero. Thus, we can reliably interpret the distance estimates as the true distance that reflects only the representational geometry and not the SNR gradient. A non-significant result compared to 0 would suggest a neural representation indistinguishable from one with full invariance.

### iEEG Experiment

#### Patients

21 epilepsy patients with implanted chronic subdural grid or strip electrodes (electrocorticography, ECoG) or stereotactic EEG (sEEG) electrodes volunteered for this study while undergoing surgical evaluation at NYU Langone Health Comprehensive Epilepsy Center (n=13) and Mount Sinai Hospital (n=8). The experiment was approved by NYU Langone Health Institutional Review Board (protocol s14-02101) and Mount Sinai Institutional Review Board (protocol 22-00529), and all patients provided written informed consent. None of these patients had a <75 Verbal Comprehension Index (VCI) score on the Wechsler Adult Intelligence Scale (WAIS-IV) or gross anatomical anomaly. Two patients were excluded due to poor task performance or recording quality (1 from NYU, 1 from Sinai). Demographic, clinical and electrode coverage information for the remaining 19 patients included in this study is available in Supplementary Table 4 and Supplementary Fig. 6.

#### Intracranial recordings

ECoG data were recorded from both electrode grids (8 × 8 contacts, 2.3 mm diameter contacts, 10 mm center-to-center spacing, Ad-Tech Medical Instrument, Racine, WI) and strips (same as grids except 1 × N contact arrangements) implanted on the cortex surface. sEEG data were recorded from electrodes inserted into the brain (8 to 12 contacts, 1.5 to 2.43 mm inter-contact distance, Ad-Tech Medical Instrument, used in 8 NYU patients and 7 Sinai patients; 5 to 18 contacts, 3.5 mm center-to-center spacing, DIXI Medical, Besançon, France, used in 4 NYU patients). The decision to implant, placement of recording electrodes, and the duration of invasive monitoring were determined solely on clinical grounds and without reference to this study.

Recordings from iEEG electrode arrays at NYU Langone Health Comprehensive Epilepsy Center were made using a NicoletOne amplifier (Natus Neurologics, Middleton, WI), bandpass filtered at 0.16–250 Hz and digitized at 512 Hz. ECoG signals were referenced to a two-contact subdural strip facing toward the skull near the craniotomy site during the recording. A similar two-contact strip screwed to the skull was used for the instrument ground.

At Mount Sinai Hospital, iEEG recordings were conducted using either XLTek monitoring units and Natus recording stations or a Neuralynx Atlas data acquisition system equipped with 196 channel low-noise electrophysiological recording capabilities. Data were sampled at with a minimum sampling rate of 1,024 Hz during recording. sEEG signals were referenced to a four-contact subgaleal strip.

#### Electrode localization

At NYU, within 24 h after surgical implantation of electrodes, patients underwent a post-operative brain MRI to confirm subdural electrode placement. Electrodes strips and grids were localized and mapped onto the pre-implant and post-implant MRI (or CT) using geometric models of the electrodes and the cortical surface^15^. First and last electrodes for depth electrodes with straight trajectories were localized manually and the remaining electrode coordinates were then automatically interpolated along the shared plane using the known inter-electrode distances. The coordinates for each electrode were then transformed into the common MNI space.

At Mount Sinai Hospital, post-operative CT images showing location of individual electrodes were co-registered with preoperative structural MRIs using normalized mutual information algorithms implemented in Fieldtrip^16^. This allowed us to determine the location of each electrode in MR-determined anatomy. Electrode anatomical location is assigned visually by contrasting electrode localizations in the co-registered CT-MR image with surgical notes and electrophysiology files. Regional localization of each contact was verified by automatic querying of anatomical Atlases (AAL, AFNI, JuBrain, and Yale Brain Atlas) and confirmed by visual examination by a neurologist or researcher.

#### Experimental Setup

Participants performed the task while reclined in their hospital beds with the use of a laptop and USB keyboard. The laptop was placed on a hospital table, with the screen ∼60cm from the participant’s eyes and in a position approximately level with their view. All images were presented with sides equaling 8.5° × 8.5° of visual angle. A photodiode was positioned over the top edge of the laptop screen and sent a signal to the amplifier when a white bar appeared simultaneously with all image presentations, allowing synchronization of image presentations and the iEEG data. Audio trigger signals, silent to the participant, were sent from the laptop’s headphone port to the amplifier to provide additional task event timing synchronization with the iEEG data. The paradigm was programmed in E-Prime from Psychology Software Tools.

#### iEEG Experimental Paradigm

The task design was adapted from previous work with Mooney images^17^ for the iEEG setting. Each trial started with a red fixation cross displayed for a pseudorandom amount of time between 1s and 2s, followed by image presentation and response collection (Supplementary Fig. 6b). Patients were instructed to fixate on the red cross whenever visible, and to respond “Yes” or “No” to the question “Can you name the object hidden in the image?” as soon as possible upon presentation of an image. “Yes” and “No” responses were collected from distinct hands on a USB keyboard. To maximize the number of trials collected within a limited amount of time, the trial concluded and the image was removed once the patient provided a response. In the absence of a response, after the 2s image presentation a response prompt was presented for up to 1s to allow additional time for responding, after which the trial ended.

Similar to earlier neuroimaging work^4, 17^, trials were grouped into blocks: three greyscale images were presented in a pseudorandom order, followed by their three corresponding Mooney images (Post images) and three novel Mooney images (Pre images) each presented twice in pseudorandom order, totaling 15 trials per block. A block was repeated to form a run of 30 trials. In total, each greyscale image was presented twice, and each Mooney image was presented four times before and four times after disambiguation. A minority of images were presented more than four times in the Pre/Post phase and more than twice for the Grayscale image due to experimenter error in a small number of recordings. At the end of each run, six verbal response trials were performed in which each Mooney image was presented for 2s and the patient was asked to verbally indicate the subject of the image if they could. The verbal responses were used to correct for any erroneous subjective recognition responses to screen out incorrect identification of Mooney images. Thus, any “Yes” keyboard responses to Mooney images during a run in which the corresponding verbal response trial was incorrect were treated as a “No” in further analyses.

17 patients completed all 14 runs; two patients completed fewer runs (12 and 13 runs each). Patients were given the opportunity for a self-directed break between runs. The runs were ordered such that the Post images and the Grayscale images in a run correspond to the Pre images from the previous run. For 14 patients, the first run consisted of only Pre trials. For the remaining five patients, in the first run, the grayscale and post-disambiguation images correspond to images used in practice trials and these trials are not used in any analysis. The entire task lasted ∼50 minutes.

### iEEG Data Analysis

*Data Preprocessing*: All sEEG data used bipolar referencing: each pair of adjacent channels were replaced by a virtual channel with location at the midpoint of the parent channels’ locations. The continuous data for each channel was imported with the MNE Python package^18^ in order to apply a notch filter to remove line noise at 60, 120, 180, and 240 Hz (zero-phase finite impulse response (FIR) filter, 3 Hz transmission bandwidth), and a high-pass filter at 0.01 Hz to remove very slow drifts. Each channels’ data was segmented according to the task runs and detrended. For each patient who had clean electrocardiogram (ECG) data, their sEEG/ECoG run data were checked for increased spectral power in the 1–1.6 Hz range and cleaned of cardiac-driven artifacts by subtracting the heartbeat evoked potential as described in previous work^19^. ECoG data is then re-referenced to the common-average reference (this was done after the earlier steps to avoid artifact from any one electrode spreading to other electrodes by contaminating the common average reference). Finally, data were manually inspected for noise (line noise, poor contact) and epileptiform activity, and contaminated segments were rejected and excluded from analysis. In addition, electrodes located outside the brain or in a lesioned area were removed prior to data preprocessing and any analysis steps.

For analysis, high gamma (50-120 Hz) power (HGP) was extracted as the amplitude envelope of the filtered data, using the Hilbert transform, and then base 10 log-transformed. The data was cut into epochs starting 1s prior to the onset of image presentation and extending to 2s after image onset for each self-report trial. Each epoch was detrended, and then baseline corrected by subtracting the mean of the 500 ms period prior to image onset. Epochs with 1,024 Hz sample rate were decimated to 512 Hz.

Only successfully disambiguated images were included in most analyses. Images were deemed successfully disambiguated when at least half the Pre trials were not recognized, at least half the Post trials were recognized, and the Grayscale image was viewed at least once (trials in which subjects were distracted away from the task were excluded). Trials presenting successfully disambiguated images are included for analysis if they are either an unrecognized Pre trial, a recognized Post trial, a recognized Grayscale trial, or the final Grayscale trial for the image. For the control analysis in Supplementary Fig. 9, recognized Pre trials were also used. Trials are clipped to end at image offset, and then smoothed with a 100-ms moving average window centered on each timepoint. Significance testing was only performed on time points at least 50 ms after image onset to avoid mixing pre- and post-stimulus signals.

### DNN Modeling — Baseline Models

We take the official code for BLT^20^ and CORnet^21^ models to train the baseline models. In order to accommodate the task which requires aggregating image features across time steps, we made minimal modifications to the architecture. The image sequence used to train baseline models is identical to those used to train our model, as described in the Method section.

The original BLT model takes the same images as the input at each time step. We change it to taking different images as the input at each time step. The model outputs log probabilities at each time step instead of only at the final time step.

The original CORnet model, specifically CORnet-S, does not have a global top-down recurrence suitable to carry information across different images. In order to make the model take image features in previous timesteps into account, we use two 1×1 convolution layers to compress the features into one channel and upsample this one-channel hidden state to match the width and height of the input image. The result is concatenated with the input image channel-wise and fed into the model in the following time step. In addition, we parametrize the initial hidden state that we concatenate with the input image in the first time step. The model outputs log probabilities at each time step instead of only at the final time step.

### DNN Modeling — Manipulation and approximations

To obtain the model’s behavioral results under grayscale image manipulations, we used the same procedure of model evaluation as in the online pilot human experiment. To apply the rotation effect and the flip effect, we apply the respective pixel level operations directly. To approximate manipulations that involve size change of the image, we apply zero padding. To approximate shift, we pad zeros to one side of the image such that the horizontal size of the new image is twice as wide. To approximate small size, we pad zeros at all boundaries of the image such that the new image is twice as tall and wide. Due to the way that the task is formulated, it is difficult to approximate enlarged images so we omit it in this analysis. The manipulated grayscale images are fed into the model as usual. The significance test is obtained with a 2-way repeated measures ANOVA with respect to the original trials or catch trials. Catch trials are implemented by replacing matching grayscale images with a completely new grayscale image that never occurs in the pre or post phase.

### DNN Modeling — Top-down conditioning signal ablation study

We linearly mix the conditioning signal with norm-matched univariate Gaussian noise with specified weight from 0 to 1, before conditioning is applied to the conditioned inference. Results were evaluated at noise weight=[0.05, 0.1, 0.2, 0.4, 0.8].

